# Integrating genomics and metabolomics to accelerate the discovery of anti-MRSA natural products from the endophytic fungus *Neocucurbitaria* sp. VM-36

**DOI:** 10.1101/2025.10.09.681339

**Authors:** Xiao Li, Rosario del Carmen Flores-Vallejo, Ting He, Jan Maarten van Dijl, Kristina Haslinger

**Affiliations:** Department of Chemical and Pharmaceutical Biology, Groningen Research Institute of Pharmacy, University of Groningen, Antonius Deusinglaan 1, 9713 AV Groningen, The Netherlands; Department of Medical Microbiology and Infection Prevention, University of Groningen, University Medical Center Groningen, Hanzeplein 1, Groningen 9700RB, The Netherlands

**Keywords:** *Neocucurbitaria* sp., whole-genome sequencing, anti-MRSA, decalin-containing tetramic acid, molecular networking

## Abstract

Endophytic fungi in medicinal plants are a rich source of bioactive natural products. Herein, we performed a comprehensive genomic and metabolic analysis of an uncharacterized endophytic fungus *Neocucurbitaria* sp. VM-36. Whole-genome sequencing and comparative analysis of the encoded biosynthetic gene clusters with six Cucurbitariaceae strains predicted its potential to produce compounds related to griseofulvin, usnic acid, hypothemycin, and phomasetin. Untargeted metabolomics confirmed several of these predictions with the presence of phomasetin analogs and isousnic acid, and uncovered a diverse range of other secondary metabolites, including specialized lipids, amino acids, and peptides, such as cyclic hexapeptides. We successfully isolated the main compound (**1**), a phomasetin analog, and show that it has bactericidal activity against different methicillin-resistant and -sensitive *Staphylococcus aureus* strains comparable in strength to vancomycin and daptomycin. Checkerboard assays with these compounds revealed mostly indifferent interactions. These findings demonstrate the antibacterial potential of compound **1** and *Neocucurbitaria* sp. VM-36.

**Graphical Abstract:** 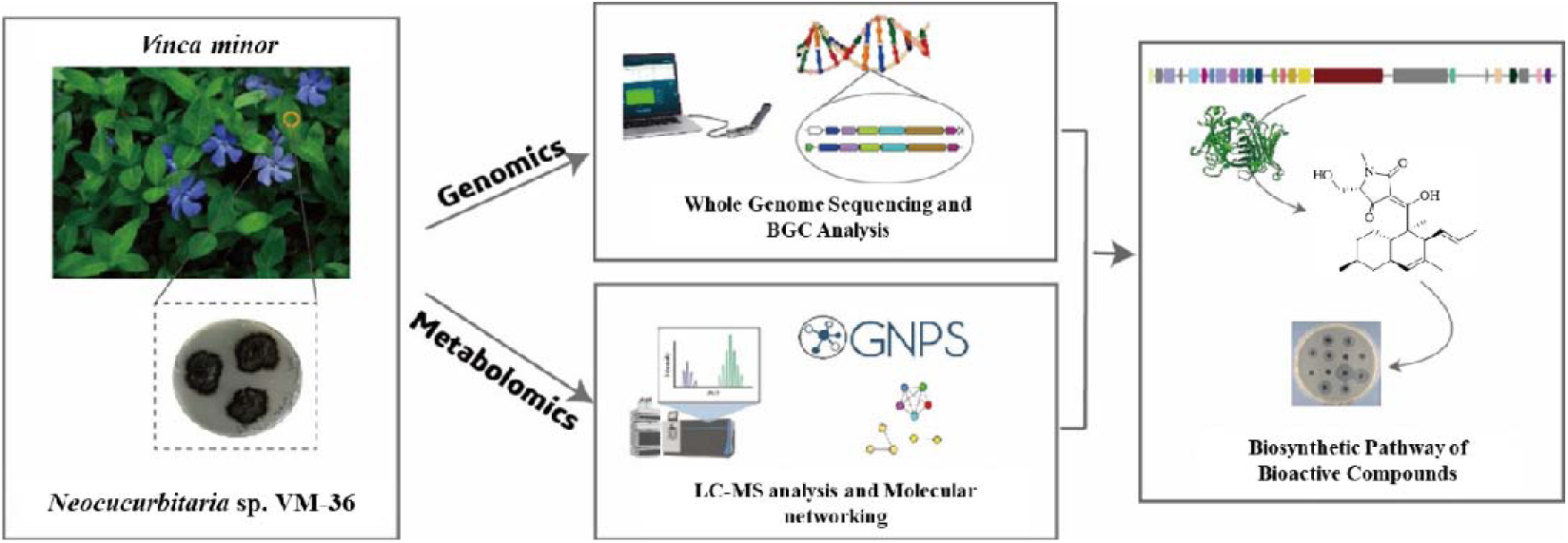

## 1. Introduction

Antibiotic resistance has become a global problem threatening human health^1^. Pathogenic bacteria can cause illnesses by producing toxins that neutralize the human immune defenses and damage healthy tissue, thereby causing harmful infections and posing a serious threat to public health^2^. While antibiotics were initially effective in treating bacterial infections, their widespread use has triggered the rapid transfer of antibiotic resistance genes between bacteria and led to prevalent bacterial resistance against clinically applied drugs^3^. For instance, certain *Staphylococcus aureus* strains, such as methicillin-resistant *S. aureus* (MRSA), have acquired the *mecA* gene, which encodes a low-affinity penicillin-binding protein (PBP2a) that allows cell wall synthesis to continue despite the presence of β-lactams^4^. Moreover, many bacterial pathogens can nowadays withstand multiple antibacterial agents, making it extremely difficult to treat the infections they cause^3^. This applies in particular to the so-called ESKAPE pathogens, which include *Enterococcus faecium*, *S. aureus*, *Klebsiella pneumoniae*, *Acinetobacter baumannii*, *Pseudomonas aeruginosa* and *Enterobacter* spp. Therefore, there is a pressing need to discover new antimicrobial agents. Endophytic fungi from medicinal plants are rich resources of novel drug leads^5^. So far, antimicrobial agents with diverse chemical scaffolds have been discovered in such endophytes, including polyketides, alkaloids, terpenoids, lactones, anthraquinones, quinones, glycosides, steroids and lignans^6^. Previously, we isolated endophytic fungi from *Vinca minor*, a medicinal plant from the *Apocynaceae* family, to obtain an in-house fungal library and conducted in-depth genomic and metabolomic analyses of two interesting fungi, namely *Fusarium* sp. VM-40^7^ and *Cosmosporella* sp. VM-42^8^. An extensive screening of our fungal library for antibacterial activity also uncovered another fascinating fungus, denoted isolate VM-36. In particular, initial disk diffusion assays showed that its crude ethyl acetate (EtOAc) extract produced a significant inhibition zone against the indicator bacterium *Bacillus subtilis* 168. Furthermore, internal transcribed spacer (ITS) sequencing analysis indicated that the fungus belongs to *Neocucurbitaria*, a little studied genus from the Cucurbitariaceae family. So far, only 20 reports on this genus can be retrieved from the PubMed database, with the earliest publication dating to 2018. Most of these publications focus on fungal ecology, with only two studies addressing fungal metabolites. In particular, a total of 15 diterpenes and their derivatives were isolated from the marine fungus *Neocucurbitaria unguis-hominis* FS685, including the neocucurbols A-H and neocucurbins A-G^9,10^. However, these compounds showed neither antibacterial activity against *Escherichia coli* and *S. aureus*, nor anticancer activity against human cancer cell lines, including the SF-268 (glioblastoma carcinoma), MCF-7 (breast cancer), HepG-2 and A549 (both liver cancer cancer) cell lines^9,10^. Altogether, the limited information that is currently available on *Neocucurbitaria* species indicates that fungi belonging to this genus still offer great opportunity for investigation.

In the present study, we conducted an in-depth analysis of the genome and metabolome of *Neocucurbitaria* sp. VM-36 to explore its capabilities for producing antibacterial secondary metabolites (SMs). We sequenced the whole genome of this fungus using Oxford Nanopore Technology, followed by contiguous assembly and annotation. We further predicted its biosynthetic gene clusters (BGCs) by AntiSMASH. We applied high-resolution, high-performance liquid chromatography-coupled tandem mass spectrometry (LC-MS/MS) and molecular networking for the analysis of its SMs. Besides, we isolated the dominant compound **1** from large-scale fermentation cultures and characterized it by 1D and 2D nuclear magnetic resonance (NMR) spectroscopy, Marfey’s method, and experimental and computational electronic circular dichroism (ECD). We also evaluated the antibacterial activity of compound **1** against several *S. aureus* strains, including interactions with vancomycin and daptomycin and potentially hemolytic effects on human red blood cells. Lastly, we propose the biosynthetic pathway of compound **1** by combining the obtained genome and metabolome data. Altogether, our study provides new insights into the biosynthetic potential of *Neocucurbitaria* sp. VM-36 and the possible development of new potent antibacterial agents for healthcare or agriculture.

## 2. Materials and Methods

### 2.1. Fungus Isolation, Morphology, and Phylogenetic Tree Analysis

The fungus *Neocucurbitaria* sp. VM-36 was isolated from healthy leaves of *Vinca minor* in November, 2021 (Groningen, The Netherlands) as previously described^7^. The strain was subsequently stored at 8 L in the department of Chemical and Pharmaceutical Biology at the University of Groningen.

To investigate the morphology of this fungus, we inoculated it onto potato dextrose agar (PDA), malt extract agar (MEA), synthetically nutrient-poor agar (SNA), and Sabouraud dextrose agar (SDA) plates. After incubation at 25 °C for 28 days, as well as 77 days for SDA, we determined the colony characteristics, including color, diameter, and mycelium morphology. Fungal hyphae were stained with lactophenol blue dye, and their microscopic features were examined with an optical microscope (Olympus BX41). Digital images were captured using a Leica camera (Heerbrugg, Switzerland) connected to the microscope.

For the molecular identification of this fungus, we compared the sequence of its Internal Transcribed Spacer (ITS) region to 18 *Neocucurbitaria* strains, with *Fusarium oxysporum* Fo47 used as the outgroup, following the method described in our recent publication^7^. We furthermore performed a multi-locus phylogenetic analysis using three barcode sequences: ITS, the RNA polymerase II subunits 2 (*rpb2*), and the large ribosomal subunit (LSU). Genome sequences of nine Cucurbitariaceae family strains, obtained from the Joint Genome Institute Genome Portal (JGI, https://genome.jgi.doe.gov/portal/) and NCBI (https://www.ncbi.nlm.nih.gov/), were included alongside *F. oxysporum* Fo47 used as an outgroup. Sequences from the three loci were concatenated and aligned using ClustalW, and the Maximum Likelihood (ML) tree was generated in MEGA (v11) with 1000 bootstrap replicates^11^.

### 2.2. Whole-Genome Sequencing, Assembly, Gene Prediction, and Annotation

DNA extraction, library preparation, sequencing, and assembly were conducted as described recently^7^. The raw reads were base-called using Guppy version 6.1.5 (Oxford Nanopore Technologies, Oxford, UK) in GPU mode using the dna_r10. 4_ e8.1_ sup. cfg model^12^. The base-called reads were subsequently filtered to a minimum length of 2 kb and a minimum quality of Q10 using NanoFilt (version 2.8.0)^13^. NanoPlot (version 1.40.0)^13^ was used to evaluate the filtered reads. Assembly was performed using Flye (version 2.9-b1778). The quality of the genome assembly was evaluated using QUAST v5.1.0rc1^14^. Bandage (version 0.8.1)^15^ was used to visualize the newly assembled genome of *Neocucurbitaria* sp. VM-36 (Figure S1). The draft assembly was subsequently polished in two rounds: first using Racon version 1.4.10 with default settings^16^ and then Medaka version 0.11.5 with default settings. The completeness of the assembly was evaluated using BUSCO 5.4.3 (ascomycota_odb10 dataset). Genome annotation was carried out using the online platform Genome Sequence Annotation Server (GenSAS, https://www.gensas.org), which provides a pipeline for whole-genome structural and functional annotation^17^. The sequencing data and genome assembly for this study were deposited in the European Nucleotide Archive at EMBL-EBI under accession number PRJEB97715. Gene Ontology (GO) annotation^18^, carbohydrate-active enzymes (CAZymes) annotation^19^, tRNA^20^ and rRNA prediction (https://github.com/tseemann/barrnap) were further performed. The biosynthetic potential of *Neocucurbitaria* sp. VM-36, along with that of six other strains with available whole genome sequences, including *Cucurbitaria berberidis* CBS 394.84, *Neocucurbitaria cava* IMI 356814, *Parafenestella ontariensis* EI-6, *Pyrenochaeta* sp. DS3sAY3a, *Pyrenochaeta inflorescentiae* CORFU0001, and *Fenestella fenestrata* ATCC 66461, was analyzed by antiSMASH fungal version 7.0.1^21^ under relaxed detection strictness and default settings.

### 2.3. Extraction of Secondary Metabolites and Molecular Networking-based LC-MS analysis

Fungal mycelium of *Neocucurbitaria* sp. VM-36 was separately transferred to two different culture media, including PDA solid medium (*ϕ* 35 mmX10 mm plate) and DPY liquid medium (20 mL medium in Erlenmeyer flask). Solid plates were incubated at 25 □ for 28 days, and their SMs were extracted with organic solvent as recently described^7^. Fungal mycelium in DPY liquid medium was cultured at 25 □ for 28 days, with shaking at 130 rpm. The fungal mycelium and broth were mixed with 20 mL EtOAc, sonicated and rotated in a rotating mixer for 1 h, respectively. The organic phase was subsequently collected and dried under a gentle stream of N_2_. The dried extract was resuspended in 1 mL of 1:1 MeOH-MilliQ water (*v*/*v*) and filtered with 0.45 µm polytetrafluoroethylene filters (Screening Devices BV company, The Netherlands).

LC-MS/MS analysis of crude EtOAc extracts was performed as described before^7^. Fresh PDA and DPY media were used as blank control groups. The acquired data was processed by Thermo Scientific FreeStyle software version 1.8. Untargeted metabolomic analysis was conducted using the online workflow on the Global Natural Products Social Molecular Networking (GNPS) website (https://gnps.ucsd.edu/) according to our previously described method^7,22^. The raw mass spectrometry data were deposited on GNPS under the accession number MassIVE ID: MSV000098902. A molecular networking analysis (https://gnps.ucsd.edu/ProteoSAFe/status.jsp?task=9099def02462433487a26b71afb487e4) was conducted using the default settings in GNPS, except that the precursor ion mass tolerance and the MS/MS fragment ion tolerance were both set to 0.02 Da. The results were visualized in Cytoscape version 3.9.1^23^. Nodes from the blank PDA and DPY media were used as background and omitted to obtain the final molecular network.

### 2.4. Fermentation, Extraction, and Isolation

The fungus *Neocucurbitaria* sp. VM-36 was grown on PDA plates at 25 □ for 28 days. 5 mycelial discs were used to inoculate a 3 □ Erlenmeyer flask containing DPY medium. 7 flasks with a total of 13.0 □ medium were cultured in a shaker at 25 □ at a speed of 130 rpm. After 30 days, the broth and mycelium were separated by filtration. The mycelium was ultrasonicated with EtOAc (3 times, 0.52 □ organic solvent each time, 1h, 30 L). The broth was extracted by EtOAc using liquid-liquid extraction (3 times, 6.5 □ solvent each time). The EtOAc extracts were combined and evaporated to dryness using a rotary evaporator (Hei-VAP Core Rotary Evaporators, Heidolph Instruments GmbH & CO. KG, Germany) to obtain a total of 1404 mg crude extract. The crude extract was further dissolved in 80% acetonitrile-water solvent and centrifuged to obtain the supernatant, which was used for MPLC separation (BUCHI Reveleris™ X2-UV System) on a C_18_ column (FlashPure EcoFlex C_18_ 12 g column from Buchi®). The analytes were eluted with solution A (water) and solution B (acetonitrile) using a step-gradient program (10 min, 0 % B; 5 min, 0%-20% B; 10 min, 20% B; 5 min, 20%-40% B; 10 min, 40% B; 5 min, 40%-60% B; 10 min, 60% B; 5 min, 60%-80% B; 10 min, 80% B; 5 min, 80%-90% B; 10 min, 90% B) to give 7 fractions. The flow rate was 10.0 mL/min. UV detection was set at 210, 230, and 294 nm. Fractions F and G were separately concentrated by a rotary evaporator under reduced pressure at 35 L, and further purified by a preparative LC (Shimadzu LC-8A preparative liquid chromatograph system, SCL-10A vp system controller, SPD-10A UV-vis detector, SIL-10AP auto injector, and FRC-10A fraction collector) on a C_18_ column (VarioPrep high-performance liquid chromatography (HPLC) Separation Column VP Nucleodur 100-5 C_18_ ec, 10 × 250 mm, 5 μm particle size) with 75% ACN isocratic elution. Column temperature was set at 40 □. The flow rate was 7 mL/min, and UV detection was set at 290 nm. 15 fractions were obtained after separation. In order to obtain pure compounds, fractions 9, 10, and 11 were purified on a Shimadzu LC-10AT high-performance liquid chromatography system equipped with a Shimadzu SPD-M20A diode array detector and eluted on a C_18_ column (EC 100/4.6 NUCLEOSHELL RP 18, 4.6 × 100 mm, 2.7 µm particle size) with isocratic 75% ACN at 0.8 mL/min flow rate, to yield compound **1** (114 mg, retention time (RT) = 8.5 min).

### 2.5. Structure Characterization

6 mg compound **1** was dissolved in DMSO-d_6_ and used for NMR analysis. NMR spectra were obtained using a Bruker Ascend Evo TM 600 NMR spectrometer (Bruker-Biospin, Billerica, MA, USA). Chemical shifts (0) are reported in parts per million (ppm) and coupling constants (*J*) are reported in hertz (Hz). MestReNova 14 was used for data analysis.

Marfey’s method was used to determine the amino acid configuration in compound **1** according to the reported method^24^ with minor modification as outlined in the Supplementary Methods.

UV and circular dichroism (CD) spectra were obtained on a JASCO V-660 spectrometer and JASCO J-815 CD spectrometer (JASCO corp., Tokyo, Japan), respectively. The wavelength scan was set to 210-400 nm. ECD calculations were performed following a reported reference^25^ with minor modifications as described below. Conformational sampling was carried out using RDKit (https://www.rdkit.org/) (ETKDGv3), and resulting conformers were energy-minimized with the MMFF94 force field. Further geometry optimization and frequency analysis were performed at the B3LYP/6-31G(d) level with the PCM solvent model for EtOH. TD-DFT calculations were performed for excited-state properties. Conformers without imaginary frequencies were subjected to ECD calculations at the B3LYP/6-31+G(d), performed without applying a solvent model due to the large computational cost and frequent convergence difficulties encountered when including solvent effects. ECD spectra were simulated using SpecDis with a half-bandwidth of 0.16–0.3 eV and averaged according to Boltzmann populations. All quantum chemical calculations were performed with Gaussian 16. Experimental and calculated spectra were compared for configuration assignment.

### 2.6. Antibacterial susceptibility assay

Several bacterial strains were used in this assay (Table S1), including the following strains from the American Type Culture Collection (ATCC) and the National Collection of Type Cultures (NCTC): *S. aureus* ATCC 29213 (methicillin sensitive strain, MSSA), *S. aureus* NCTC 8325 (MSSA), *Enterococcus faecalis* ATCC 29212, *Enterobacter cloacae* ATCC 13047, *Klebsiella pneumoniae* ATCC 13883, *Acinetobacter baumannii* ATCC 17978, *Pseudomonas aeruginosa* ATCC 27853, and *Escherichia coli* ATCC 25922. In addition, we used *S. aureus* strain Newman ^26^, *S. aureus* NE1688 (Sle1) (MRSA) from the Nebraska transposon library^27^, which carries a transposon insertion in the *sle1* gene ^28^, and the two clinical MRSA isolates D15 and D17, which were respectively associated with community- and hospital-acquired infections^29^. The minimum inhibitory concentrations (MICs) and minimum bactericidal concentrations (MBCs) of compound **1** were determined using the broth microdilution method in 96-well microtiter plates as described in the Supplementary Methods. The lowest concentration of compound **1** that completely inhibited bacterial growth was determined as the MIC, and the MBC was determined as the lowest concentration of compound **1** that reduced the viable bacterial count on agar plates by 99.9%. Vancomycin and daptomycin were used as positive controls. Bacteria treated with the respective vehicles were used as negative control.

Bacterial growth kinetics of compound **1**, vancomycin, and daptomycin against 1 MSSA and 4 MRSA strains at final concentrations of 2, 4, and 8 µg/mL were monitored by measuring the optical density at 600 nm (OD_600_) values. Bacterial viability before compound addition and at the endpoint was determined by colony-forming unit (CFU) enumeration, as detailed in the Supplementary Methods. All experiments were performed in triplicate.

The drug interaction of compound **1** with daptomycin or vancomycin against the three MRSA strains, USA300, D15 and D17 was determined using the checkerboard assay in 96 well microplates as described in the Supplementary Methods.

### 2.7. *In vitro* cytotoxicity on human erythrocytes

To evaluate the effect of compound **1** on human erythrocytes, a hemolysis assay was performed as detailed in the Supplementary Methods. Briefly, compound **1** was dissolved in DMSO at concentrations of 200, 100, 10, and 1 μg/mL. DMSO at 1%, 0.5%, 0.05% and 0.005% v/v concentrations were used as blanks. Sodium dodecyl sulfate at a final concentration of 0.1% w/v was used as the positive control and Dulbecco’s phosphate-buffered saline (DPBS) pH 7.2 served as a negative control. Vancomycin and daptomycin were included at the same final concentrations as compound **1**. Double-distilled water and NaCl 0.9% w/v were evaluated as their blanks, respectively. All groups were tested in triplicates.

### 2.8. Medical ethics committee approval

Blood donations from healthy volunteers were collected based on written informed consent with approval of the medical ethics committee of the University Medical Center Groningen (UMCG; approval no. Metc2012-375), and in accordance with the Declaration of Helsinki guidelines.

## 3. Results

### 3.1. Morphological Characterization of *Neocucurbitaria* sp. VM-36

The *Neocucurbitaria* sp. VM-36 was cultured on different nutrient media and the macroscopic features of its colonies were recorded (Figure 1A). PDA and MEA media appeared to be more favorable for mycelial growth and conidia production compared with SNA and SDA media. On PDA plates, the colonies presented a diameter of 27-37 mm after 28 days with black mycelium and a textured center. On MEA plates, the colonies reached a diameter of 35-40 mm with a brown and white floccose mycelium and a non-hardened center. The fungus formed smaller colonies on SNA plates, with a diameter of 15-25 mm and white velvety aerial hyphae. Colonies on SDA plates were only 12-20 mm in diameter and had a markedly different yellow color. They appeared like solid particles at first and only at the final stage of the 28-day cultivation, a gray-white mycelium emerged on the top. Accordingly, we continued to culture the fungus on SDA plates until day 77 at which time point the colonies reached a diameter of 28-37 mm with brown mycelium and a textured center. On the reverse side the colonies were brown-yellow in the center and black in the periphery.

**Figure 1.**
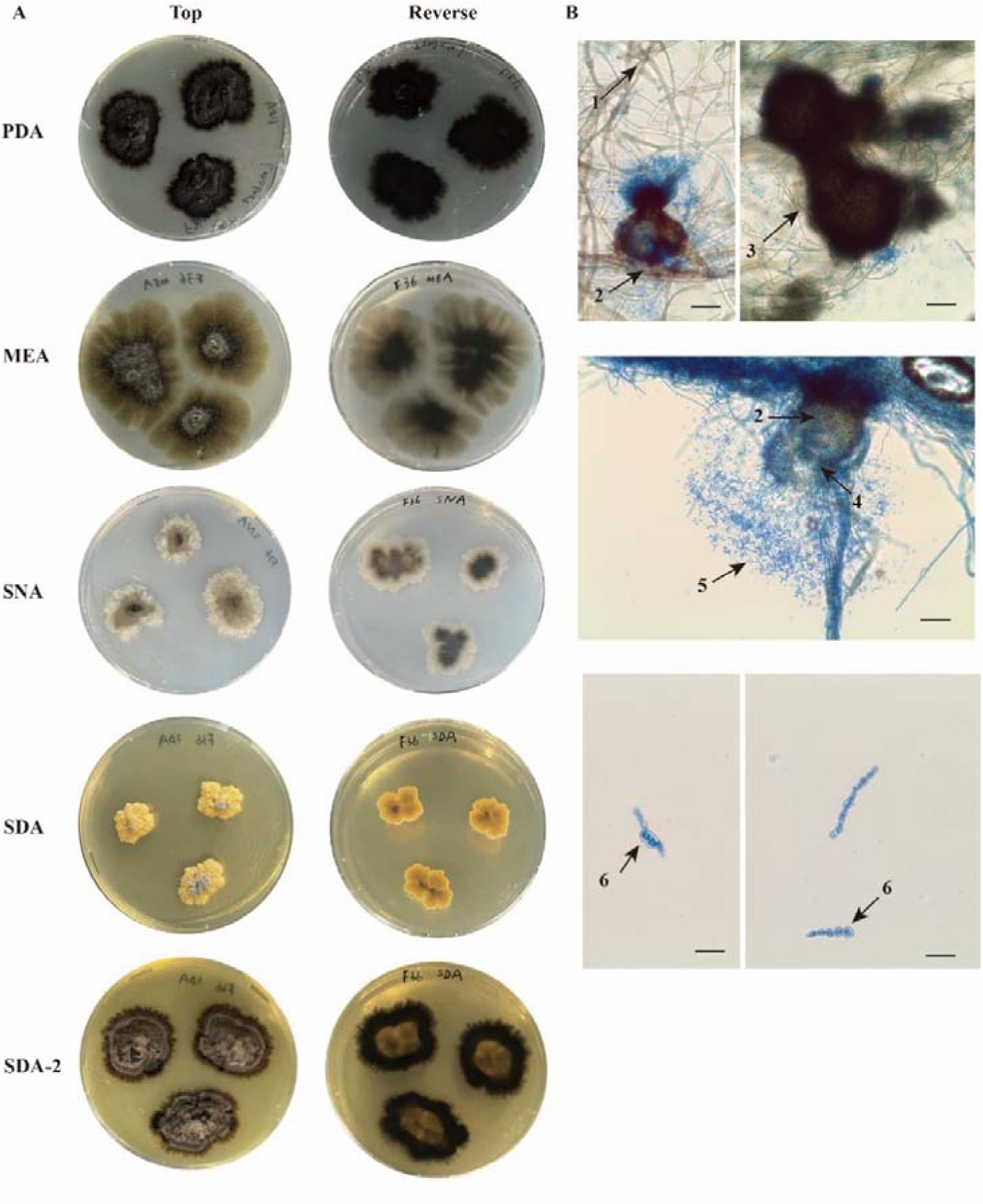
Macroscopic (A) and microscopic (B) characteristics of *Neocucurbitaria* sp. VM-36 on different culture media (PDA, MEA, SDA, and SNA), incubated at 25 °C for 28 days or 77 days (SDA-2). (1) Hyphae; (2) Pycnidium-Conidiomata; (3) Pycnidia; (4) Conidiophores; (5) Conidia; and (6) Chlamydospore chains formed on SDA medium. Scale bars = 50 µm.

Microscopy revealed that the fungus formed spherical pycnidia, an asexual structure lined with conidiophores for producing asexual conidiospores (Figure 1B). In addition, *Neocucurbitaria* sp. VM-36 also produces chlamydospores when grown on SDA plates, which support the survival under adverse conditions due to the formation of particularly thick walls.

### 3.2. Genomic analysis

To characterize the genetic make-up of *Neocucurbitaria* sp. VM-36, we sequenced, assembled, annotated, and analyzed its whole genome (Table S2). With the highly complete (97.2 % according to BUSCO) and contiguous assembly with only 19 contigs at hand, we performed additional phylogenetic analyses to more accurately locate our isolate in the *Neocucurbitaria* genus. Both single-locus and multi-locus analyses consistently showed that *Neocucurbitaria* sp. VM-36 is closely related to *N. salicis-albae* CBS 144611 (Figure 2), a species previously isolated from a *Salix alba* twig in Germany^30^.

**Figure 2.**
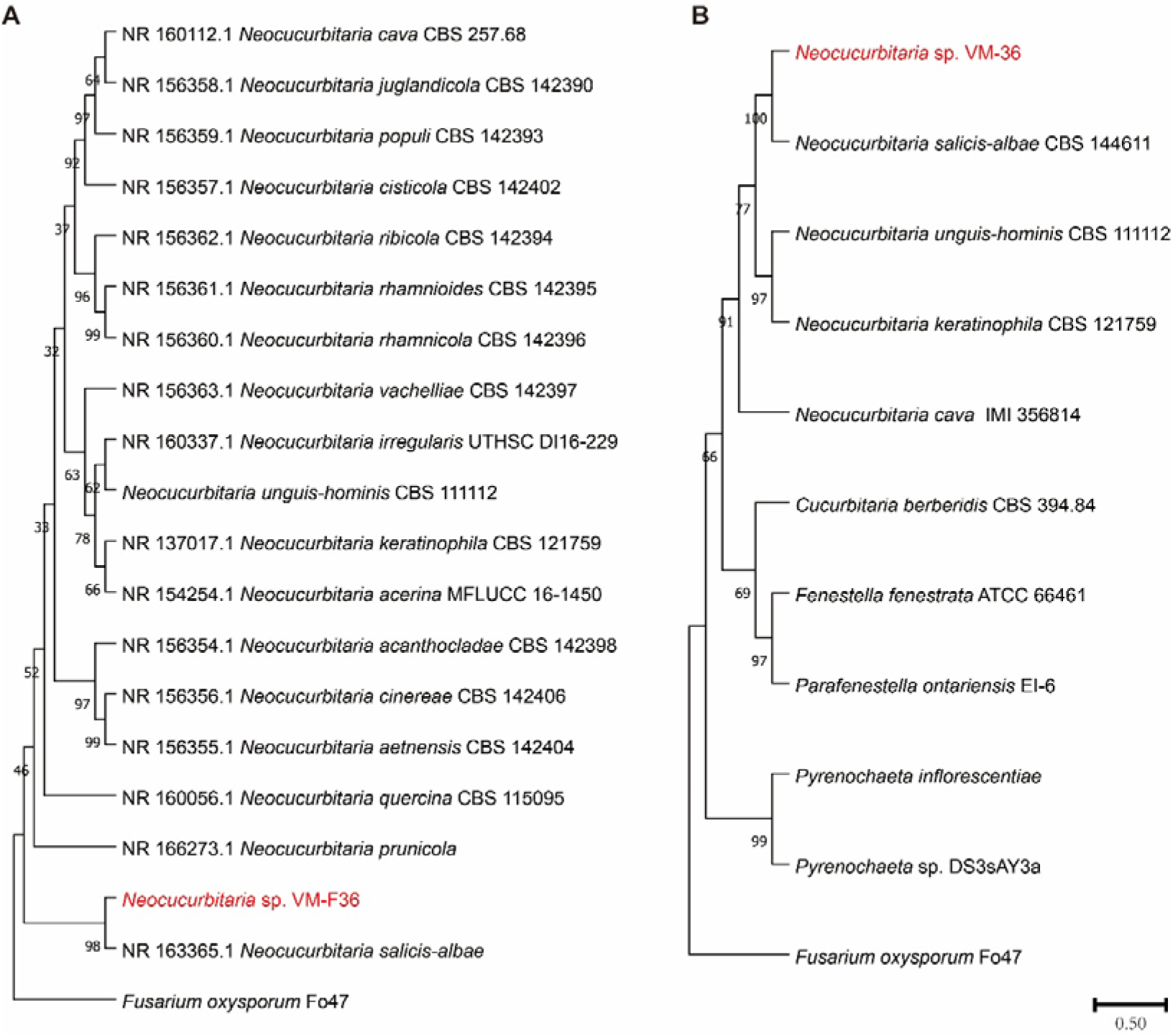
Phylogenetic analysis of *Neocucurbitaria* sp. VM-36. (A) ML phylogram of *Neocucurbitaria* sp. VM-36 based on the ITS sequence. (B) Maximum Likelihood (ML) phylogenetic tree based on concatenated nucleotide sequences of *rpb2*, ITS, and LSU. *F. oxysporum* Fo47 was used as an outgroup in both analyses.

Functional annotation of the assembly revealed that the genome encodes 11,817 proteins, 29 rRNAs and 120 tRNAs. Based on Gene Ontology (GO) analysis the top 50 terms are categorized into biological processes (24%), molecular functions (52%), and cellular components (24 %) (Figure S2A). Among these, carbohydrate metabolism was prominently represented, with 224 genes annotated under “carbohydrate metabolic process,” 694 under “hydrolase activity,” and 129 under “pectate lyase activity”. More in-depth analysis revealed the presence of 571 CAZyme-related genes. This highlights the ecological role and biotechnological potential of *Neocucurbitaria* sp. VM-36 in plant biomass decomposition and carbohydrate metabolism. This potential appears to be shared with other Cucurbitariaceae species (Figure S2B), suggesting conserved metabolic strategies likely shaped by shared ecological niches and evolutionary pressures.

To also explore the biosynthetic potential of *Neocucurbitaria* sp. VM-36, we predicted the presence of BGCs in its genome and compared them to the predicted BGCs of six strains in the Cucurbitariaceae family. A total of 34 putative BGCs were identified in *Neocucurbitaria* sp. VM-36, a number comparable to the average BGC count (36) predicted across other Cucurbitariaceae fungi (Figure 3A, Table S3). These BGCs were categorized into five major types: 10 polyketide synthase (PKS)-type BGCs, 9 ribosomally synthesized and post-translationally modified peptide (RiPP)-like BGCs, 9 non-ribosomal peptide synthetase (NRPS) and NRPS-like BGCs, 3 terpene-type BGCs, and 3 hybrid BGCs (2 NRPS+T1PKS and 1 T1PKS+ fungal-RiPP-like). Annotation against the MIBiG database revealed that 11 of the 34 BGCs were similar to known biosynthetic pathways (Figure 3B, Figure S3). Notably, BGCs associated with choline and metachelin C production were conserved across all seven analyzed fungi, suggesting a common biosynthetic capability within the family.

**Figure 3.**
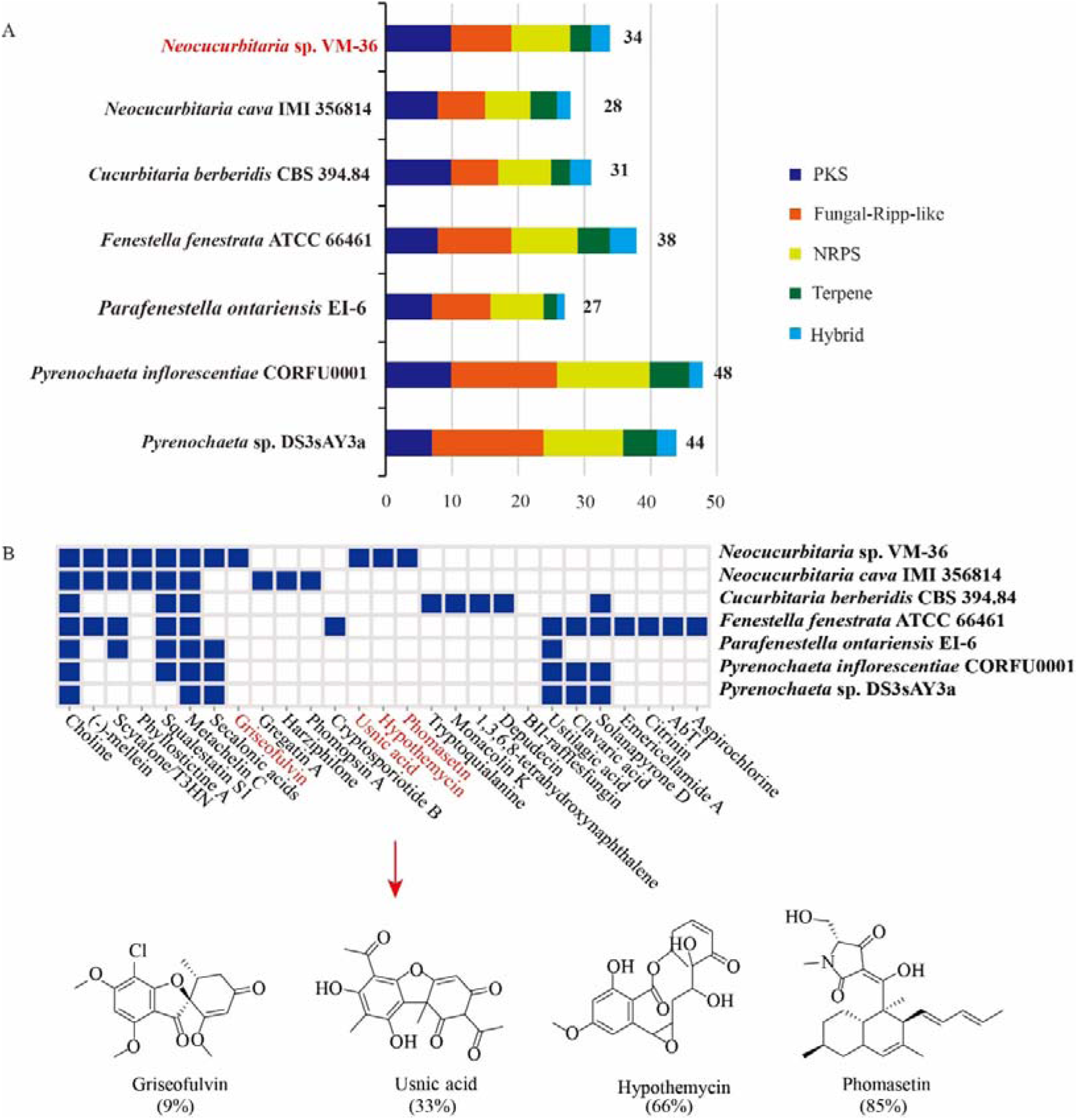
Comparative analysis of predicted BGCs in 7 strains of the Cucurbitariaceae family. (A) Number of BGCs in each genome by BGC type; (B) Presence (blue) or absence (white) of known BGCs in the 7 strains and chemical structures associated with 4 of the known BGCs which were only identified in *Neocucurbitaria* sp. VM-36.

Additionally, BGCs responsible for the biosynthesis of squalestatin S1, secalonic acids, scytalone/T3HN, (-)-mellein, and phyllostictin A were identified not only in *Neocucurbitaria* sp. VM-36 but also in select species within the Cucurbitariaceae family. Importantly, four BGCs, putatively involved in the production of compounds related to hypothemycin, usnic acid, griseofulvin, and phomasetin, were unique to *Neocucurbitaria* sp. VM-36. These unique clusters suggest a distinct biosynthetic repertoire, potentially contributing to novel bioactivities (Supplementary Note S1).

### 3.3. Molecular Networking-Based Secondary Metabolite Identification

So far, only 15 compounds have been isolated from *Neocucurbitaria* strains, encompassing macrocyclic and phomactin diterpenes and their derivates^9,10^, and a total of 28 compounds were isolated from the Cucurbitariaceae family (Table S4, Figure S4). Such few reports on SMs from this family aroused our interests for exploring the chemical compounds formed by *Neocucurbitaria* sp. VM-36 and their biosynthetic origin in this fungus. Thus, we performed an untargeted metabolomic analysis of fungal crude extracts from PDA and DPY medium, compared to medium blank controls. The final molecular network consists of 1521 nodes and 1896 edges (Figure 4, Table S5 and Figure S5). 19 compounds were putatively identified via library matching with existing MS2 databases and 7 compounds were inferred from matched adjacent nodes (Figure 4 and 5).

**Figure 4.**
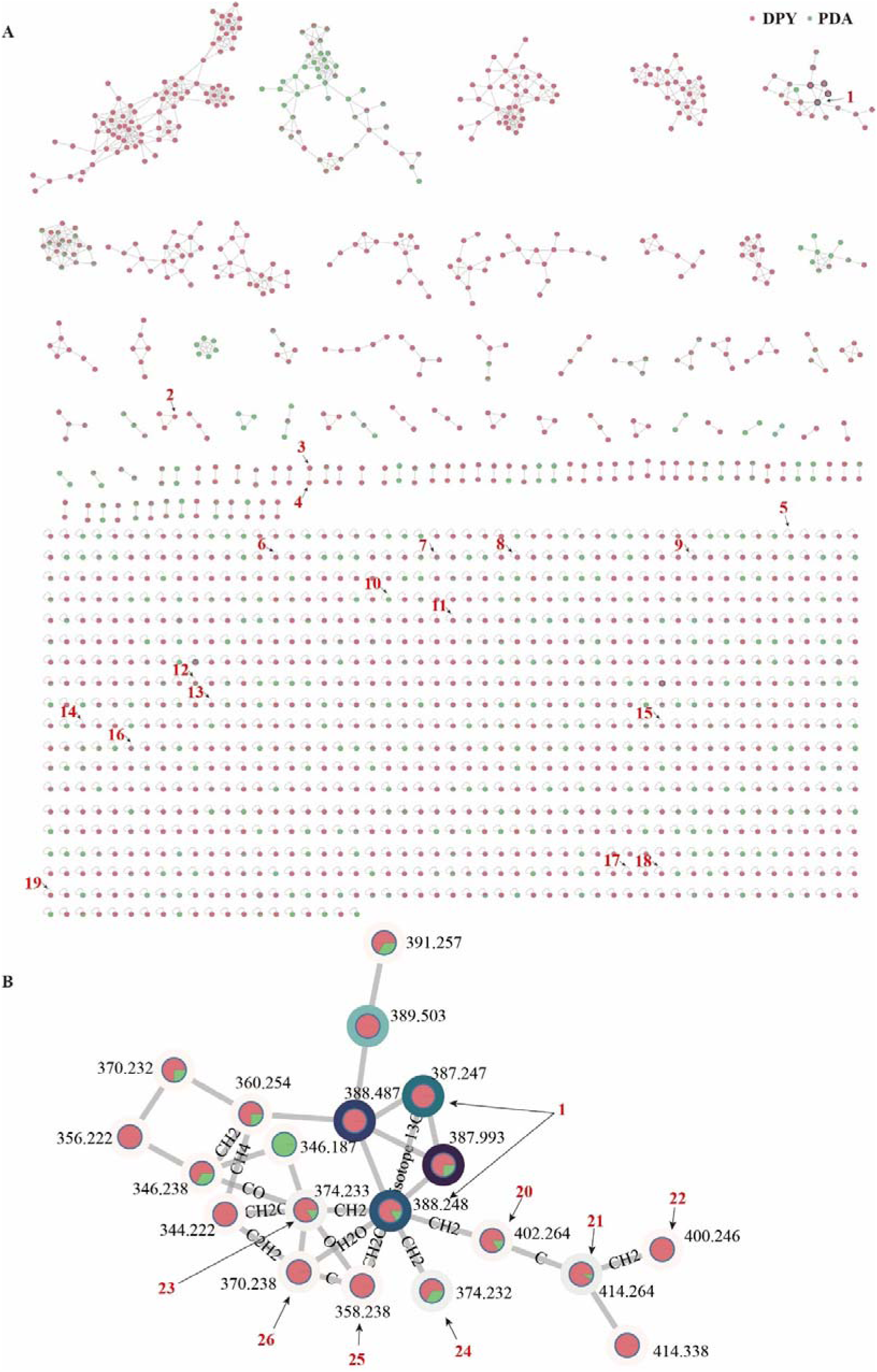
Molecular network of SMs in crude EtOAc extracts of *Neocucurbitaria* sp. VM-36 grown in PDA solid medium and DPY liquid medium. (A) Overview of the molecular network, with annotated SMs; (B) Molecular network of tetramic acid components containing decalin rings (cluster 5). Node colors represent signals in samples from PDA solid medium (green) and DPY liquid medium (red). Darker shades of node borders indicate higher metabolite abundance in samples based on peak area integration of the base peak, and purple represents the highest abundance. Edges represent the structural similarity between nodes.

**Figure 5.**
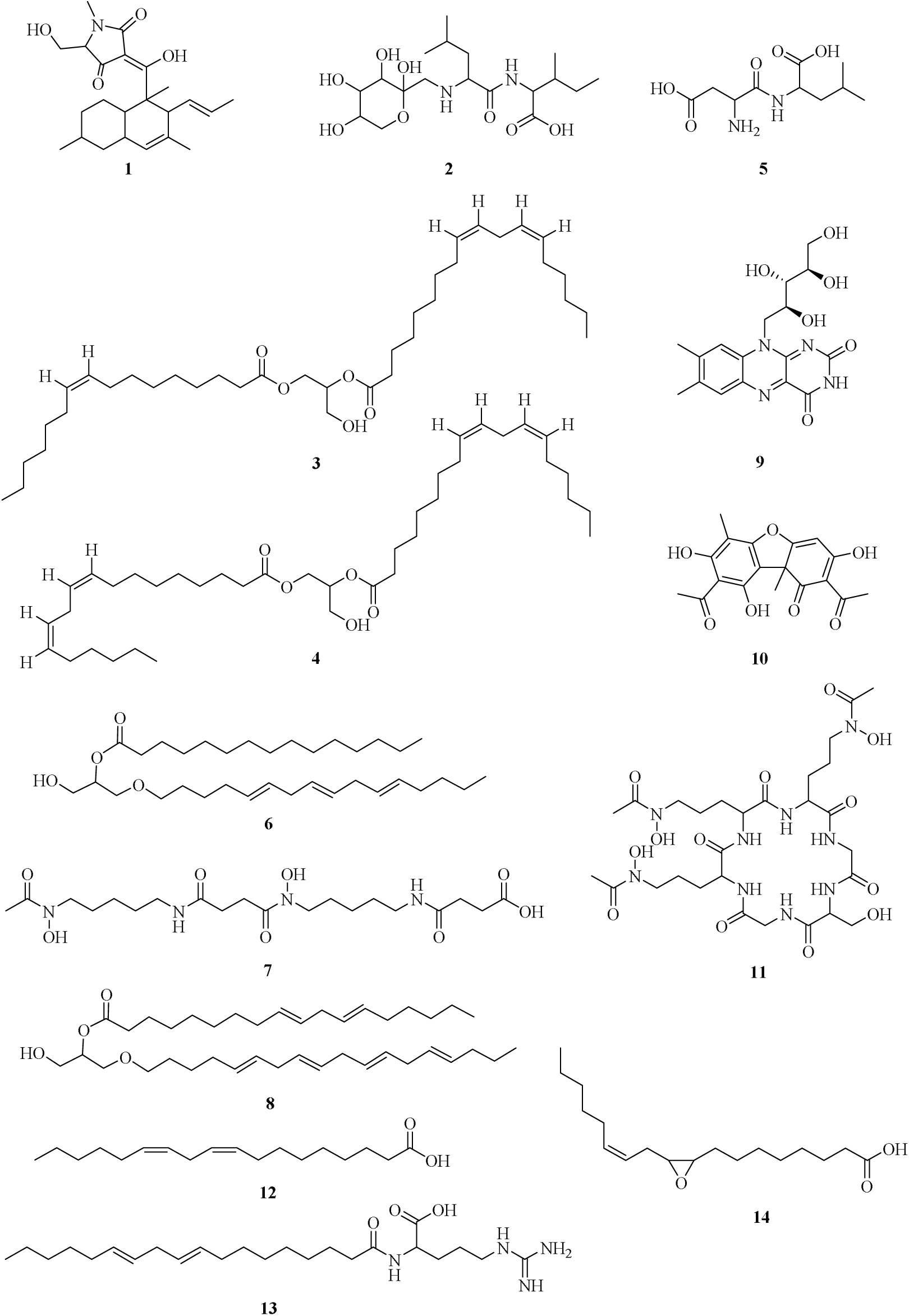
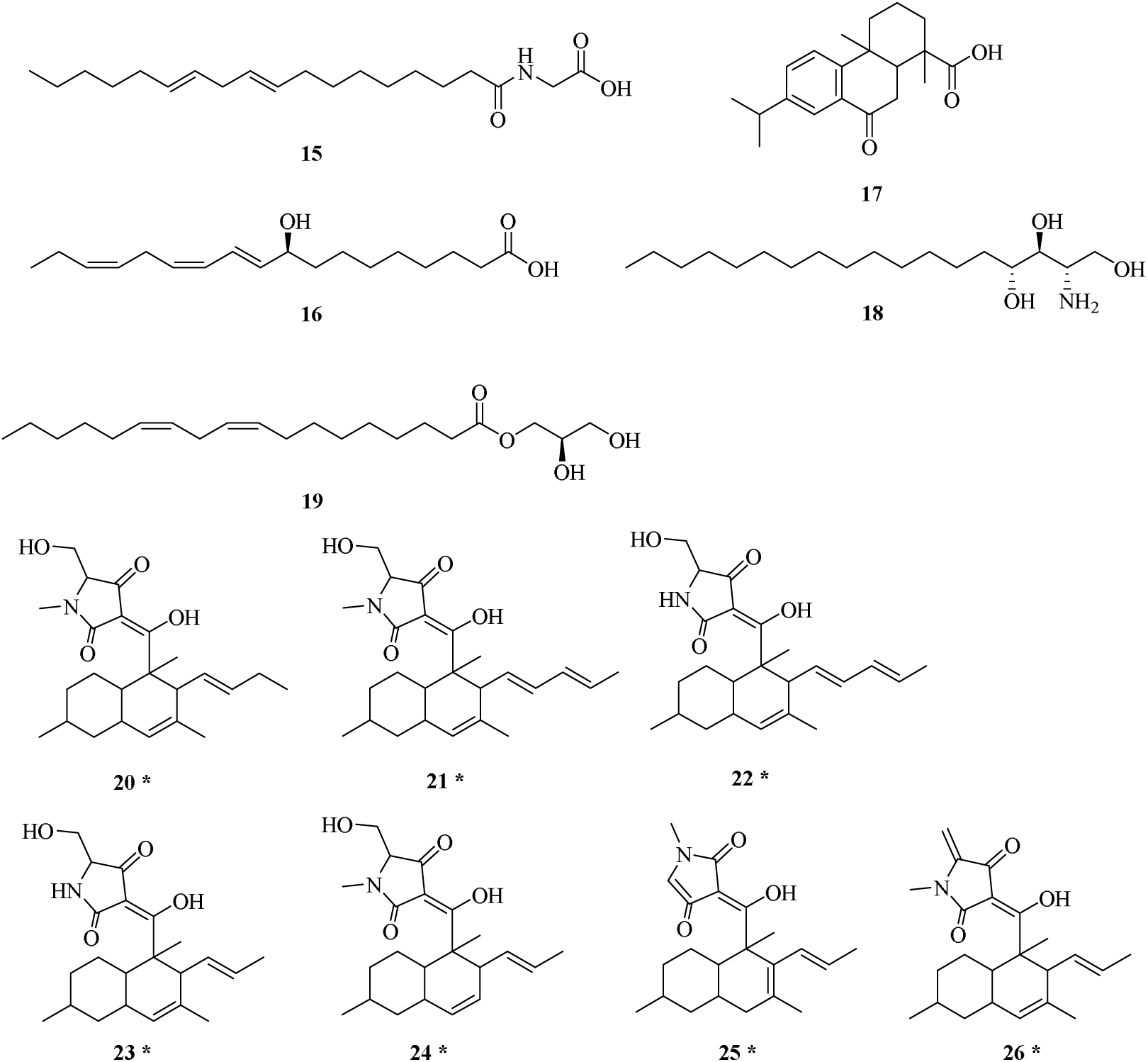
Chemical structures of annotated compounds putatively identified in the crude extracts of *Neocucurbitaria* sp. VM-36 grown in PDA solid medium and DPY liquid medium (* indicates putative structures according to the molecular network).

The major constituents with high abundance in the crude extracts are annotated to be CJ-21058 analogues in cluster 5 (Figure 4B), which are tetramic acid-type components containing decalin rings or DTAs (decalin-containing tetramic acids). Based on the MS2 fragments (Table S5), we inferred 7 additional tetramic acid-type compounds (Supplementary Note S2).

Among the less abundant compounds, a total of 9 lipids, including compounds **3** (DG(16:1/18:2/0:0)), **4** (DG(18:2/0:0/18:2)), **6** (AEG(o-16:3/15:0)), **8** (AEG(o-18:4/18:2)), **12** (linoleic acid), **14** (9(10)-EpOME), **16** (9S-Hydroxy-10E,12Z,15Z-octadecatrienoic acid), **18** (phytosphingosine), and **19** (1-linoleoylglycerol), are annotated as singleton nodes in the molecular network. Six amino acid-containing or peptide compounds, including compounds **2** (FruLeuIle), **5** (Asp-Leu), **7** (4-[5-[[4-[5-[acetyl(hydroxy)amino]pentylamino]-4-oxobutanoyl]-hydroxyamino] pentylamino]-4-oxobutanoic acid), **11** (N-[3-[5,8-bis[3-[acetyl(hydroxy)amino]propyl]-14-(hydroxymethyl)-3,6,9,12,15,18-hexaoxo-1,4, 7,10,13,16-hexazacyclooctadec-2-yl]propyl]-N-hydroxyacetamide/ desferriferricrocin), **13** (Arg-C18:2), and **15** (Gly-C18:2), are also found scattered across the network. Compound **11** is a cyclic hexa-peptide hydroxamate siderophore, with the general formula RC(O)N(OH)RL (*N*-hydroxy amides)^31^. Compounds of this type do not only help fungi to chelate iron for survival under iron-restricted conditions, but they have also attracted attention due to their medicinal value as potent antibacterial or antitumor agents^31^. In *Neocucurbitaria* sp. VM-36, we hypothesize that the BGC encoded in region 8.2 (hybrid BGC of NRP-metallophore and NRPS) is likely to be responsible for producing **11**. Although there are no results with Known-Cluster-Blast, the core NRPS gene in the BGC shows amino acid similarity with an epichloenin A synthetase

(MIBIG protein-AET13875.1, BGC0001250), with 26% identity and 82.7% coverage. Its enzymatic product, epichloenin A, is an extracellular siderophore related to ferrirubin, which is required to maintain the mutualistic interaction of the endophyte *Epichloe festucae* with the perennial ryegrass *Lolium perenne*^32^. Specifically, secretion of epichloenin A enables the symbiotic fungus to compete for plant iron while avoiding fungal over-growth inside the host plant^32^. Moreover, another core gene in the NRP-metallophore BGC of *Neocucurbitaria* sp. VM-36 shares similarity with an L-ornithine_N-monooxygenase (MIBiG protein CAJ96465.1) from BGC0000330, with 42% amino acid identity and 87% coverage, suggesting that it is probably responsible for forming an N-hydroxylated and N-acylated ornithine moiety in **11**^33^.

Node **10** is predicted to be the [M + H + CH_3_OH]^+^ adduct of isousnic acid, an isomer of usnic acid. The cluster in region 20.2 in *Neocucurbitaria* sp. VM-36 has a Known-Cluster-Blast hit with BGC0002483, producing usnic acid, and 33% of its encoded enzymes show amino acid similarity with the query sequence. In particular, it encodes enzymes similar to the non-reducing PKS (methylphloracetophenone synthase) and a cytochrome p450 (methylphloracetophenone oxidase), which are responsible for the production of usnic acid^34^. Usnic acid, a toxic dibenzofuran, has been discovered in lichens and was extensively reported to have a variety of pharmacological effects, including remarkable antimicrobial, antitumor, antiviral, and antiparasitic activities^35,36^. It is active against Gram-positive bacteria, such as *B. subtilis* and *S. aureus*, by inhibiting RNA and DNA synthesis and blocking DNA replication and elongation, but it is not active against Gram-negative bacteria^36^. Although usnic and isousnic acid are regioisomers, the latter is comparatively poorly investigated, with only a few studies reporting on its antimicrobial and anti-inflammatory properties^37,38^.

Overall, many clusters in the molecular network, especially with three or more nodes, could not be assigned to compound classes or specific compounds. They are potentially new compounds and need to be further explored by compound isolation and structural identification.

### 3.4. Structural Characterization of Compound 1

Compound **1** (Figure 6) is the most abundant constituent of *Neocucurbitaria* sp. VM-36. It is predicted to have a tetramic acid-containing decalin scaffold based on its MS2 fragmentation. We purified it from large-scale fungal culture as a white amorphous powder. Its molecular formula was established to be C_23_H_33_NO_4_ based on its *m/z* 388.2489 value ([M + H]^+^ ion, calculated 388.2488), with 8 degrees of unsaturation. The absorbance spectrum of compound **1** in EtOH with maxima at A_max_ (£) 252 (4.05) and 291.8 (4.08) nm (Figure 6C) is similar to the structurally related compound phomasetin^39^. Furthermore, the absolute configuration of the serine-derived moiety in compound **1** is also □ (Figure S6A), as known for phomasetin^39^.

**Figure 6.**
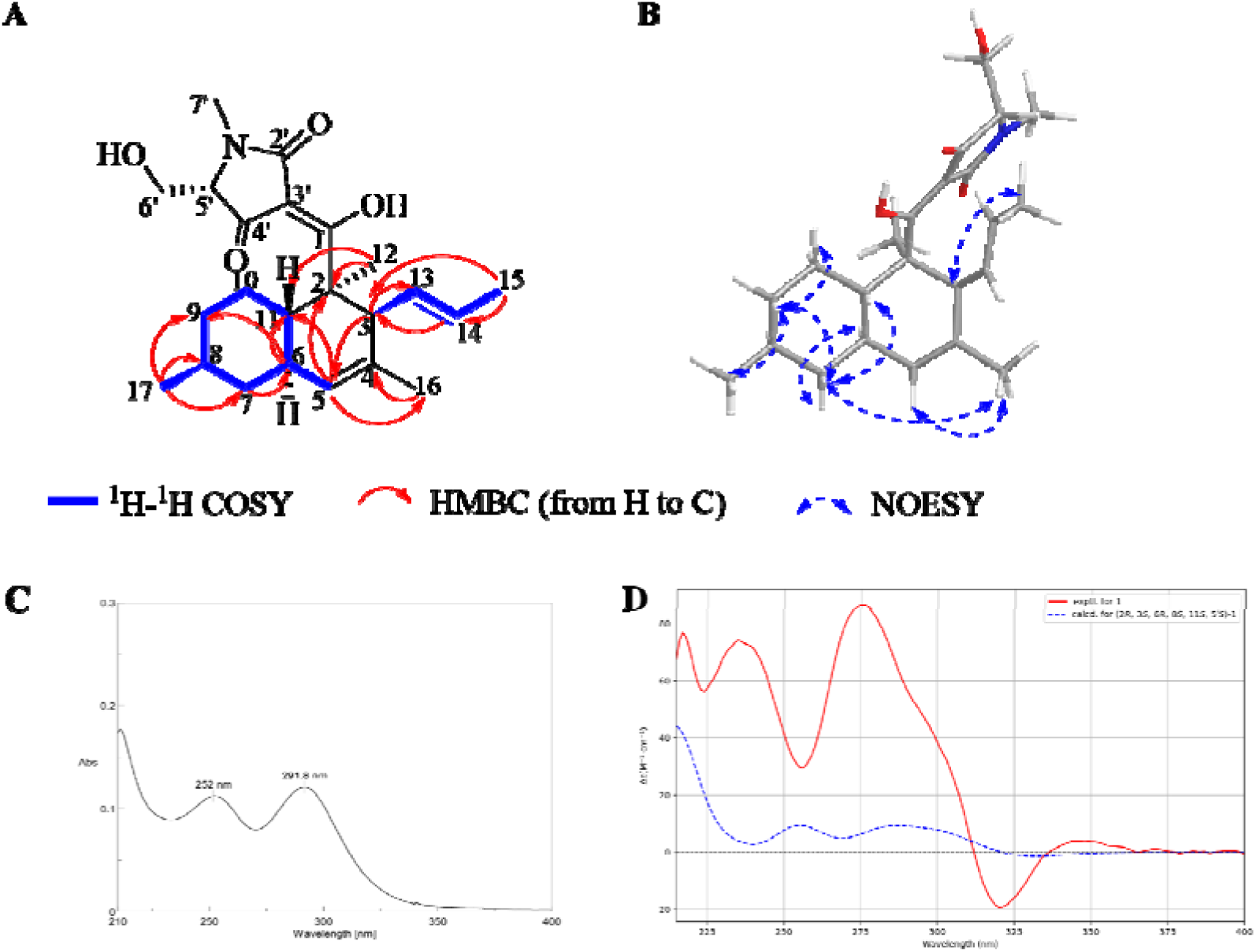
Physico-chemical characterization of compound **1**. Chemical structure with key two-dimensional NMR correlations (COSY and HMBC (A) and NOESY (B)); UV spectrum (C) and experimental and calculated ECD spectra (D).

Next, we determined the 3D structure of compound **1** by NMR^24^ and ECD. According to the 1D NMR (Table 1 and Figures S7-S8) and MS2 data (Table S5), compound **1** has one -C_2_H_2_ unit less and two sp^2^-carbon signals less than phomasetin, indicating the absence of one double bond in compound **1** compared to phomasetin^40,41^.

**Table 1.** ^1^H NMR (600 MHz) and ^13^C NMR (150 MHz) data of compounds **1** (in DMSO-d6)

| Position | Compound **1** |  |
| --- | --- | --- |
| | $\delta_{\text{H}}$, Multi. ($J$ in Hz) | $\delta_{\text{C}}$, Type |
| 1 | - | 203.14, C |
| 2 | - | 49.13, C |
| 3 | 4.02, m | 48.06, CH |
| 4 | - | 132.00, C |
| 5 | 5.16, s | 126.05, CH |
| 6 | 1.78, m | 39.11, CH |
| 7 | 0.81, m | 42.50, $\text{CH}_2$ |
|  | 1.75, m |  |
| 8 | 1.47, m | 33.30, CH |
| 9 | 1.01, m | 35.80, $\text{CH}_2$ |
|  | 1.70, m |  |
| 10 | 0.98, m | 27.98, $\text{CH}_2$ |
|  | 1.90, d (10.9) |  |
| 11 | 1.57, t (10.05) | 39.59, CH |
| 12 | 1.31, s | 13.79, $\text{CH}_3$ |
| 13 | 4.92, m | 132.99, CH |
| 14 | 5.09, m | 131.29, CH |
| 15 | 1.49, d (5.7) | 17.98, $\text{CH}_3$ |
| 16 | 1.52, s | 22.47, $\text{CH}_3$ |
| 17 | 0.88, s | 22.82, $\text{CH}_3$ |
| 2' | - | 170.82, C |
| 3' | - | 107.16, C |
| 4' | - | 194.78, C |
| 5' | 3.42, m | 65.08, CH |
| 6' | 3.79, dd (11.8, 2.9) | 58.63, $\text{CH}_2$ |
|  | 3.66, m |  |
| 7' | 2.92, s | 27.17, $\text{CH}_3$ |

This feature matches that of CJ-21,058 and its conformational isomers, the hyalodendrins A and B^41,42^. Compared to the CJ-21,058 NMR data^41,42^, compound **1** has similar signals except for slight differences in the chemical shifts (Table 1)^41,42^. Most notably, the ^13^C chemical shift for C-1 in compound **1** is 203.14 ppm instead of 199 ppm, for C-2’ 170.82 ppm instead of 177 ppm, for C-3’ 107.16 ppm instead of 100 ppm, and for C-4’ 194.78 ppm instead of 190 ppm^41^.

Also, the 2D NMR correlations (^1^H-^1^H COSY and ^1^H-^13^C HMBC) of compound **1** indicate high structural similarity with the hyalodendrins A and B^41^ (Figure 6A and S9-S12). ^1^H-^1^H NOESY correlations (Figure 6B), including H6/H7a, H7b/H9a, H7b/H11, H9a/H10b, and H9a/H17, indicate that proton H-6 is on the opposite face of H-11 and H17, forming a trans-decalin structure.

To determine the absolute configuration of **1** and to pinpoint the functional groups at the C-1 and C-2’ positions, we collected experimental ECD spectra and compared them to calculated ECD spectra (Figure 6D and S6B). The experimental ECD spectrum indicates the presence of a mixture of two isomers, with different spectral contributions across wavelength regions. In the 210-225 nm range, the experimental peak shape aligns well with both computed spectra, suggesting overlapping contributions. From 225-250 nm, the experimental peak closely matches that of isomer B (Figure S6B) in both position and shape, though with higher intensity, while isomer A (Figure 6D) shows a weaker, red-shifted response. Within the 270-300 nm region, the experimental ECD spectrum displays a pronounced positive peak. A similar feature is present in the computed spectrum of isomer A (Figure 6D), albeit with reduced intensity. In contrast, isomer B (Figure S6B) shows only a weak signal, remaining near the baseline. Beyond 300 nm, both the experimental and calculated spectra exhibit a negative peak. However, the intensity is significantly stronger in the experimental spectrum. The lack of solvent treatment in the final calculations, due to the high computational cost and frequent convergence difficulties, combined with limited conformer sampling and a restricted number of excited states considered, likely contributes to the discrepancies observed between experiment and theory. Overall, the data support a mixed-isomer system with varying contributions across the ECD spectrum. Based on this analysis, we assigned the (2*R*, 3*S*, 6*R*, 8*S*, 11*S*, 5’*S*) configuration and propose that interconversion occurs at the C-1 and C-2’ positions in solution, resulting in the isolated compound existing as a mixture of the two tautomers shown in Figures 6A and S6B.

### 3.5. Proposed Biosynthetic Pathway for Compound 1

The cluster in region 13.1, classified as a hybrid T1PKS/NRPS BGC, shares partial similarity with the phomasetin biosynthetic gene (*phm*) cluster from *Pyrenochaetopsis* sp. RK10-F058 (BGC0001738) and is therefore likely to give rise to compound **1** (Figure 7). In particular, the core enzyme NeoA shares 72% amino acid identity with the PKS-NRPS hybrid Phm1 in the *phm* cluster^43^. It comprises one highly-reducing PKS module and one NRPS module with a terminal reductase (TD) domain. The PKS module is likely complemented by a stand-alone enoyl reductase (NeoB), thus producing the linear polyene intermediate via 7 cycles of polyketide chain elongation, reduction, and methylation. Subsequently, L-serine is incorporated by the NRPS module and upon release from the assembly line, the tetramic acid moiety is formed by the Diekmann cyclase function of the terminal reductase (TD) domain.

**Figure 7.**
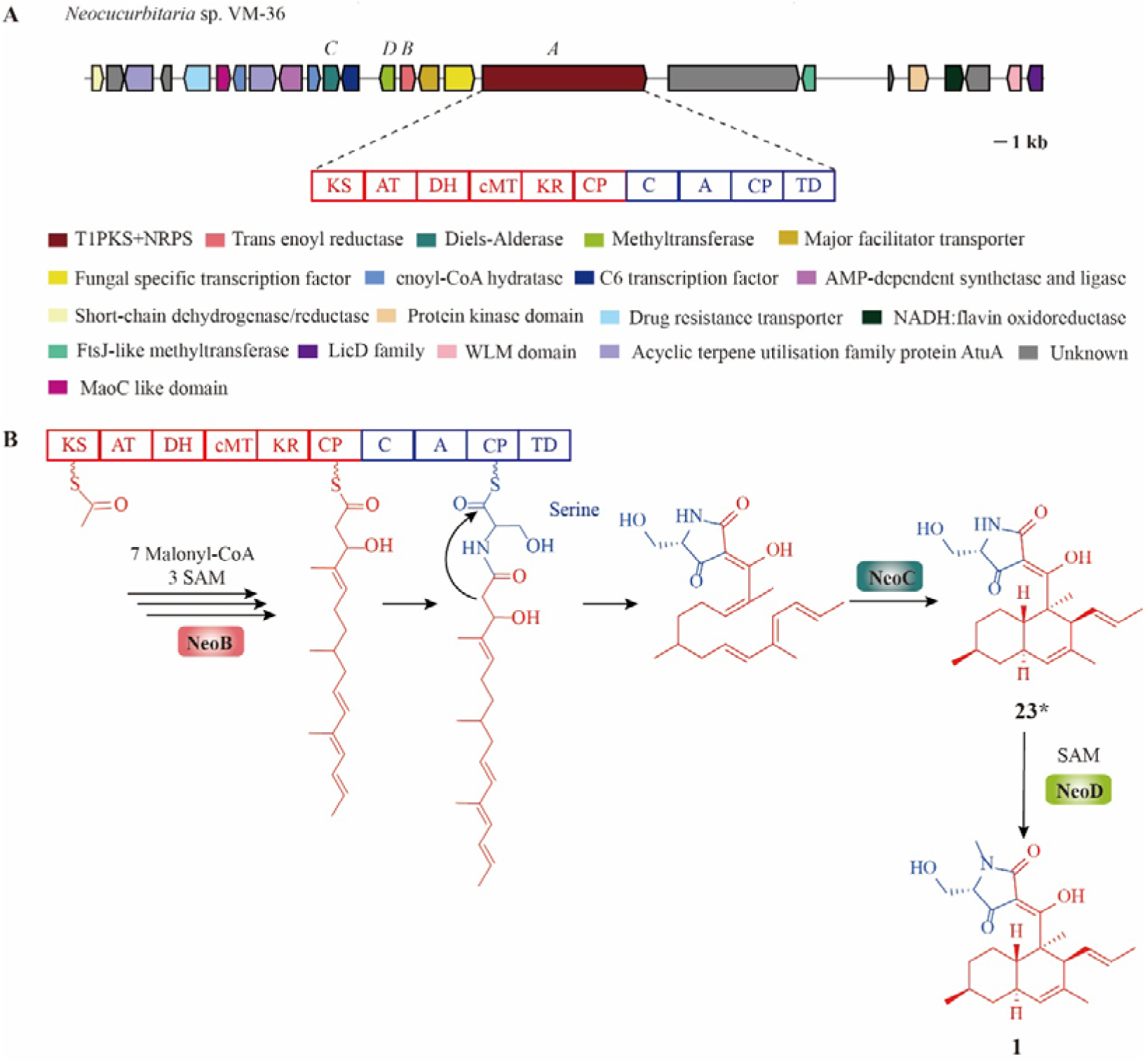
Gene cluster (A) and proposed biosynthetic pathway (B) of compound **1**. β-ketosynthase (KS), acyl-transferase (AT), dehydratase (DH), carbon-methyltransferase (cMT), β-ketoreductase (KR), and acyl-carrier protein (ACP), condensation (C), adenylation (A), peptidyl carrier protein (PCP), and terminal reductase (TD).

NeoC, displaying 57% amino acid identity with Phm7 in the *phm* cluster^43–45^, is predicted to be a Diels-Alderase (DAse)^40–42^. DAses can catalyze intramolecular Diels-Alder cycloadditions between a conjugated diene and substituted alkene via C-C bond formation, yielding a cyclohexene with up to four chiral centers^44^. Evolutionary analysis of NeoC with some representative [4+2]-cyclases shows that NeoC and Phm7 are in the same clade, with a close relationship to homologs gNR600, PvhB, and Fsa2 (Figure 8). Although DAses can theoretically form four stereoisomers of *trans-* and *cis-*decalin, most of these enzymes, including Phm7, Fsa2, and gNR600, tend to stereo-selectively produce the *trans-*decalin structure in conjunction with a fungal PKS^43–48^. PvhB, involved in the biosynthesis of varicidin A in *Penicillium variabile*, is an exception of a fungal DAse to form cis-decalin, most likely driven by the presence of an electron-withdrawing carboxylate on the diene, which kinetically favors the exo-transition state in the DA reaction^46,49^. Among the trans-decalins, equisetin and phomasetin produced by Fsa2 and Phm7, respectively, represent the two possible enantiomers with (2*S*, 3*R*, 8*S*, 11*R*) and (2*R*, 3*S*, 8*R*, 11*S*) configurations, respectively^44,45^. Computational modeling, including molecular dynamics simulations and quantum chemical calculations, demonstrated that the reactions proceed through synergetic conformational constraints assuring transition state-like substrate folds and their stabilization by specific protein-substrate interactions^44^. Furthermore, the flexibility of bound substrates is largely different in two enzymes, suggesting the distinct mechanism of dynamic regulation behind these stereoselective reactions^44^.

**Figure 8.**
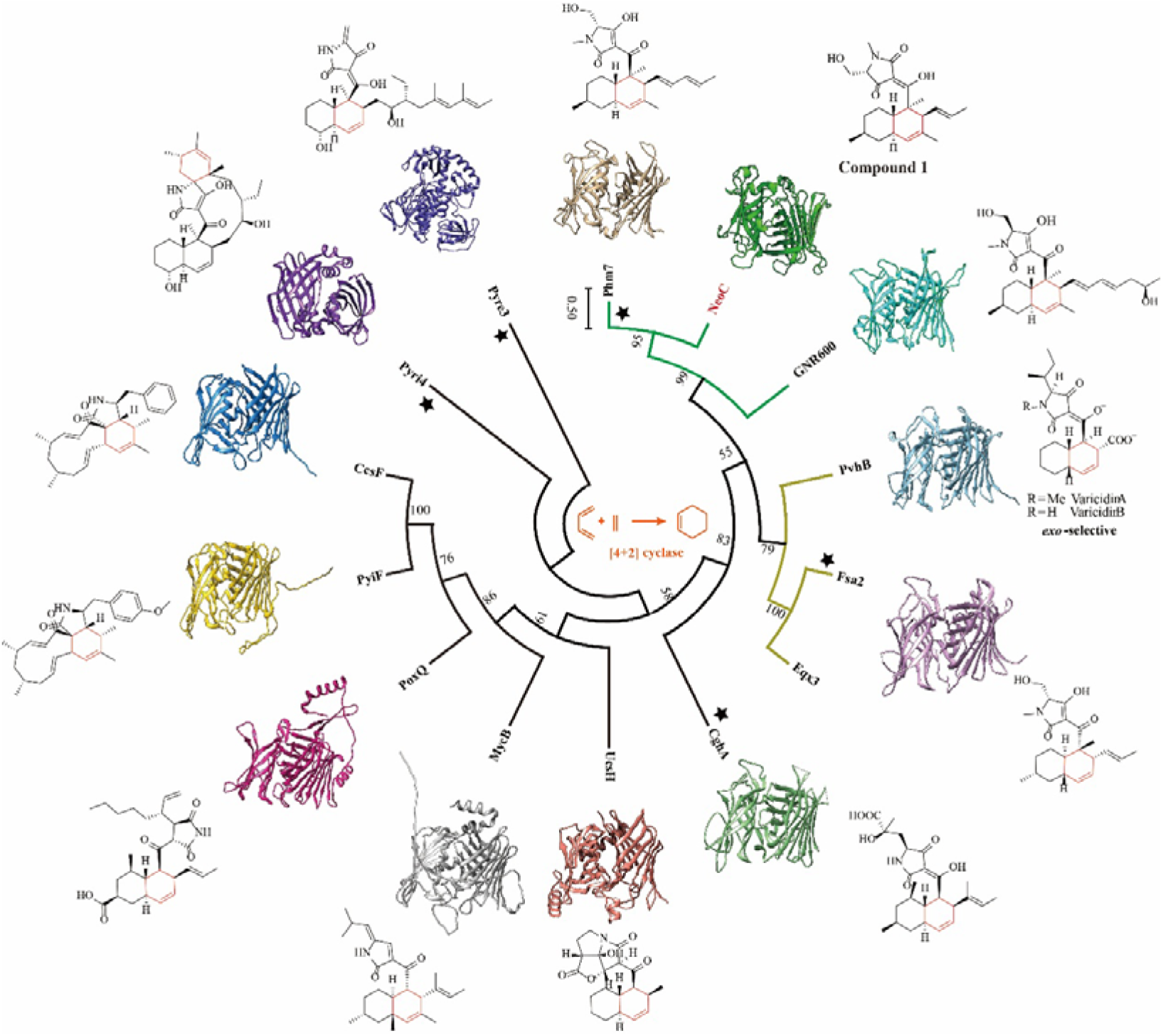
Evolutionary analysis of NeoC with other decalin-forming [4+2]-cyclases with Pyrl4 as an outgroup. Enzymes marked with stars have reported crystal structures, and the remaining protein structures were predicted by AlphaFold3. The substructure resulting from the enzyme-catalyzed cyclo-addition is highlighted in red in the respective compound. Enzyme selection and figure layout were adapted from Xu and Yang (2021)^50^.

The dominant compound in the crude EtOAc extract of *Neocucurbitaria* sp. VM-36, compound **1,** has a trans-decalin scaffold with the same configuration as phomasetin, but its side chain at position C3 is shorter than that of phomasetin. We also detected trace amounts of phomasetin itself (compound **21**) and other structurally related compounds (**20, 22**-**26**) in our untargeted metabolomics analysis, indicating that the same BGC may give rise to a group of related compounds with compound **1** being the favored one under the chosen culture conditions. This suggests that the PKS-NRPS domain has some flexibility in the number of polyketide chain elongation cycles with a preference for the compound **1** precursor and/or that the preference of the DA enzyme for the compound **1** precursor leads to the dominance of compound **1**.

When comparing the amino acid sequences of NeoC and Phm7 (Figure S13), we note that most of the active amino acid residues are conserved^44,45^, except for residue 258, where isoleucine replaces threonine. Whether this amino acid change influences the substrate or product scope of the enzyme, remains to be investigated.

Lastly, NeoD, annotated as methyltransferase, shares 71% amino acid identity with the methyltransferase Phm5 in the *phm* cluster. We hypothesize that the -NH group in the tetramic acid moiety of an N-demethylated precursor (putatively compound **23** in the molecular network) can be methylated by NeoD to generate the N-methylated product compound **1**. In addition to the enzymes described above, there are still many other functional enzymes encoded in this gene cluster. Whether they contribute to further structural complexity remains to be explored.

### 3.6. Bioactivity Evaluation of Compound 1

As the crude extract of *Neocucurbitaria* sp VM-36 exerted a strong antibacterial effect, we were curious to see whether compound **1** was responsible for or contributed to this effect. Thus, we screened the compound against ESKAPE pathogens and found that it exerts strong inhibitory effects against several Gram-positive bacterial strains, including *E. faecalis* ATCC 29212 (MIC = 4 µg/mL), *S. aureus* ATCC 29213, *S. aureus* NCTC 8325, *S. aureus* Newman (MIC = 1 µg/mL), and *S. aureus* HG001 (MIC = 2 µg/mL), while it was inactive against the Gram-negative bacterial strains (MICs > 64 µg/mL) (Table S6). We further determined the MIC and MBC values of compound **1** against four MRSA strains, including *S. aureus* USA300, two clinical isolates *S. aureus* D15 and D17, and the transposon mutant *S. aureus* NE1688 (Sle1), as well as the MSSA reference strain *S. aureus* ATCC 29213 (Table 2). Compound **1** shows strong antibacterial activity against these MRSA and MSSA strains (MIC = 0.5 - 1 µg/mL), similar to the positive controls vancomycin (MIC = 1 µg/mL) and daptomycin (MIC = 0.5 - 2 µg/mL). The MBC values of compound **1** (2 - 4 µg/mL) were slightly higher than those of vancomycin and daptomycin (MBC = 1 - 2 µg/mL). With MBC/ MIC ratios ≤ 4, we conclude that compound **1** shows bactericidal effects on *S. aureus* USA300, D15 and D17, and ATCC 29213. Strain *S. aureus* NE1688 (Sle1) appears to be more sensitive to growth inhibition by low concentrations of compound **1** but it is similarly sensitive to lethal doses as the other strains.

**Table 2.** Antibacterial activity evaluation of compound **1** against *S. aureus*.

| Strain | Compound **1** |  |  | Vancomycin |  | Daptomycin |  |
| --- | --- | --- | --- | --- | --- | --- | --- |
|  | MIC<br>(µg/mL) | MBC<br>(µg/mL) | MBC/<br>MIC<br>ratio | MIC<br>(µg/mL) | MBC<br>(µg/mL) | MIC<br>(µg/mL) | MBC<br>(µg/mL) |
| *S. aureus* ATCC 29213 (MSSA) | 1 | 2 | 2 | 1 | 1 | 0.5 | 1 |
| *S. aureus* USA300 (MRSA) | 1 | 4 | 4 | 1 | 1 | 1 | 1 |
| *S. aureus* clinical isolate D15 (MRSA) | 1 | 2 | 2 | 1 | 2 | 1 | 2 |
| *S. aureus* clinical isolate D17 (MRSA) | 1 | 4 | 4 | 1 | 1 | 1 | 1 |
| *S. aureus* USA 300 NE1688 (Sle1) (MRSA) | 0.5 | 4 | 8 | 1 | 2 | 2 | 2 |

We also analysed the bacterial growth curves of the strains in the presence of compound **1** at MBC levels (2, 4, and 8 µg/mL) to monitor the dose-dependent change of optical density (Figure 9). After the exposure to compound **1**, the bacteria stopped growing and the OD_600_ values declined. Also, the maximum OD_600_ values of all bacterial cultures were significantly reduced in the presence of compound **1** compared to the vehicle controls (DMSO). This shows that compound **1** not only has a dose-dependent bactericidal activity but also that it induces bacterial lysis, with the strongest effects observed at 8 µg/mL. To further corroborate this observation, we assessed the bacterial viability before and after 40 h treatment by counting CFUs per mL liquid culture (Figure S14). Compared with the positive controls, compound **1** exerts stronger bactericidal activity against *S. aureus* ATCC 29213, USA300, D15 and D17 at 8 µg/mL and 4 µg/mL, with little to no bacterial colonies observed. Against *S. aureus* NE1688 (Sle1), compound **1** showed greater bactericidal activity than both controls at 8 µg/mL, but was less effective than daptomycin at 4 µg/mL. As expected, at the lowest tested concentration (2 µg/mL), none of the compounds exerts a significant bactericidal effect.

**Figure 9.**
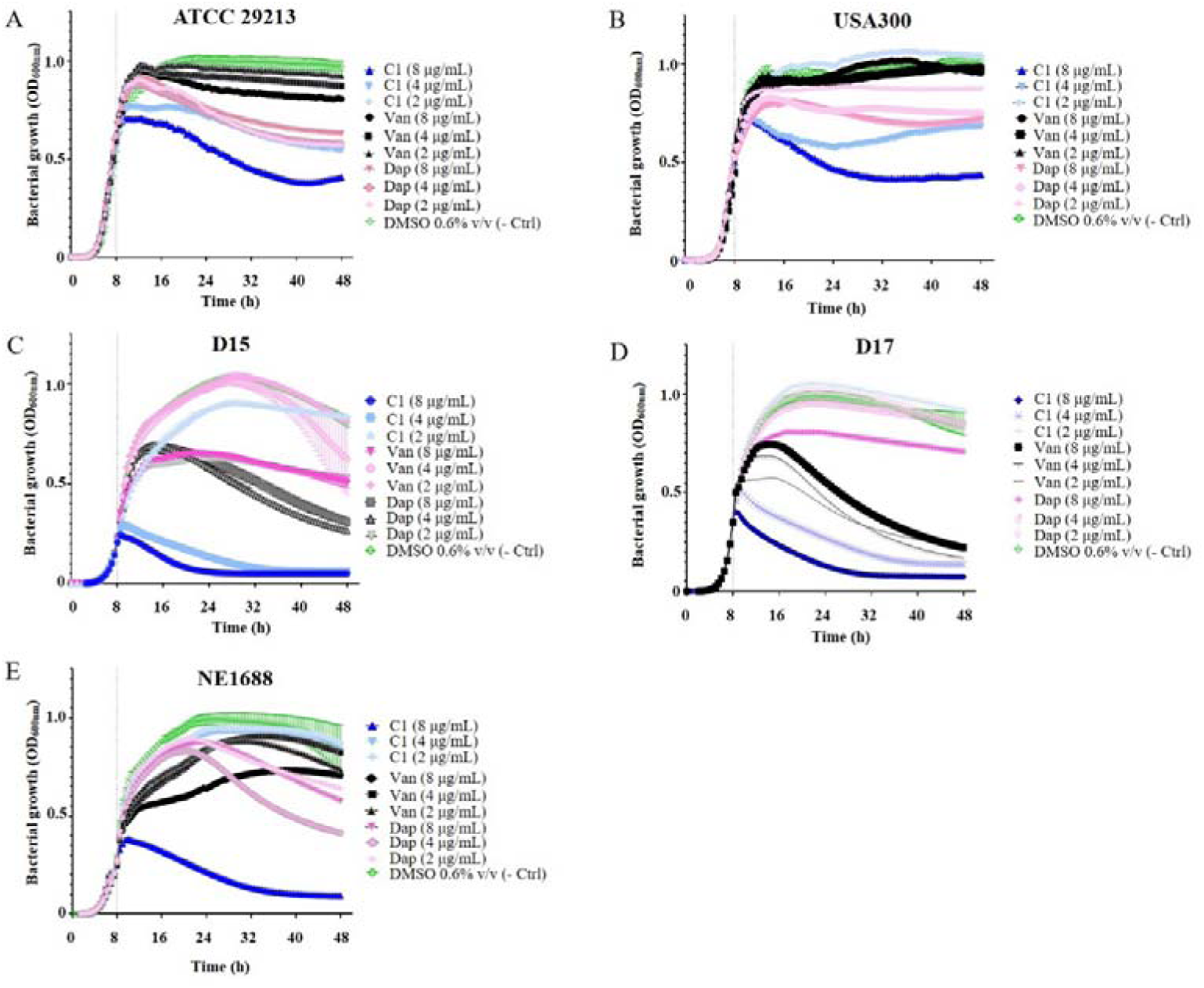
Growth curves of 5 *S. aureus* strains in the presence of compound **1**, or the positive controls vancomycin and daptomycin at 2, 4 and 8 µg/mL. (A) *S. aureus* ATCC 29213 (MSSA); (B) *S. aureus* USA300 (MRSA); (C) *S. aureus* D15 clinical isolate (MRSA); (D) *S. aureus* D17 clinical isolate (MRSA); (E) *S. aureus* NE1688 (MRSA). The scattered line (---) in the x-axis indicates the time point at which the compound was added (t = 8 h). Results are expressed as mean OD_600nm_ values and error bars represent standard deviation (n = 3).

Lastly, we also assessed whether a combination treatment of three MRSA strains (*S. aureus* ATCC 29213, D15, and D17) with compound **1** and daptomycin or vancomycin had any synergistic or antagonistic effects (Figure 11). Among 22 treatment combinations in the checkerboard assay, none exhibited antagonistic effects. Most interactions were indifferent, with a minimum fractional inhibitory concentration index (FICImin) of 1.25, except for the combination of compound **1** and daptomycin, which showed an additive effect against the clinical isolate D17 from a hospital-acquired infection (FICImin = 0.625). These findings indicate that compound **1** holds potential for combination therapy with established antibiotics, particularly daptomycin, to improve efficacy against particular MRSA strains, as exemplified with the D17 strain. The lack of antagonistic interactions suggests a favourable safety profile for such combination treatments.

**Figure 11.**
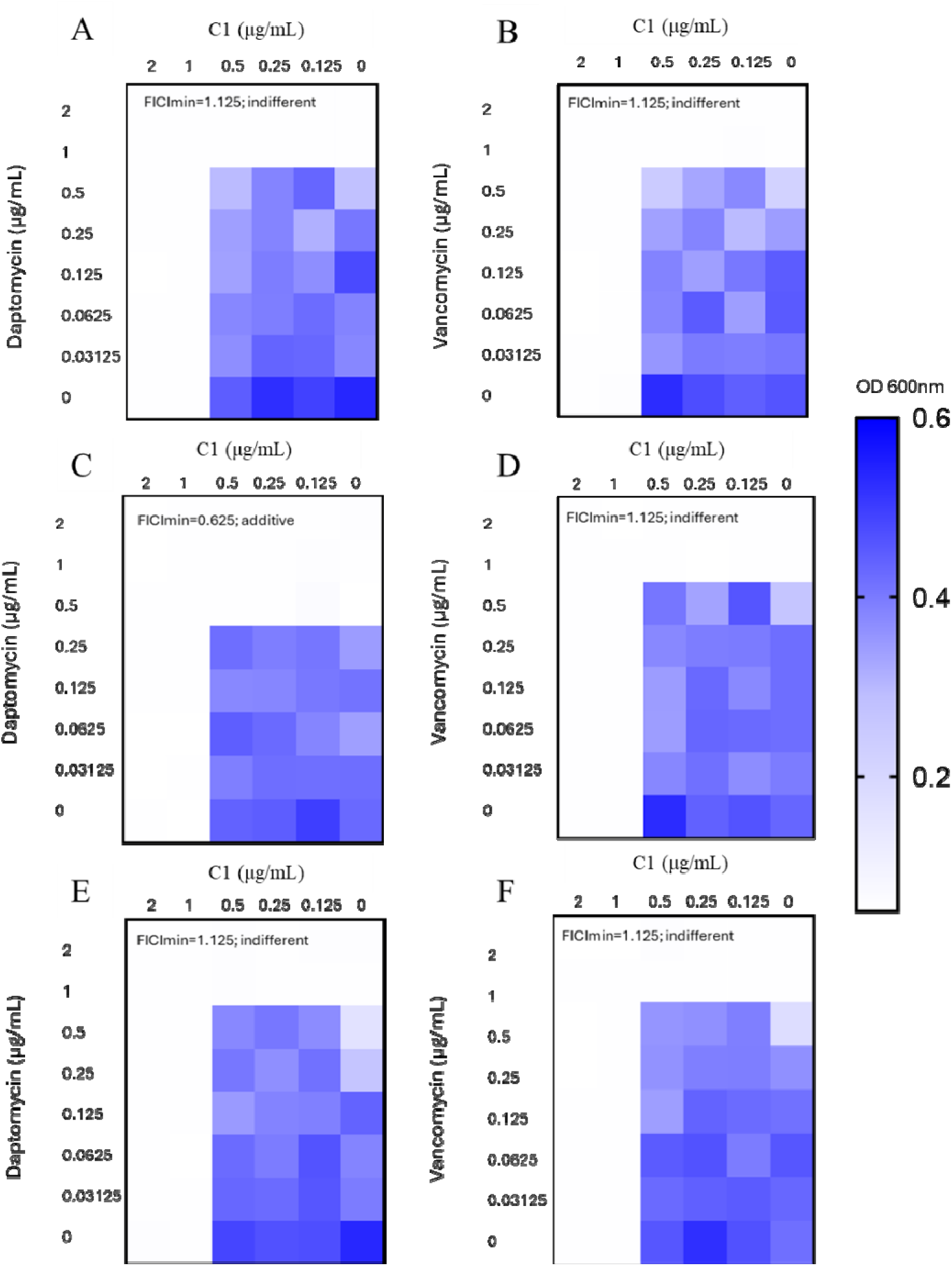
Effects of combination treatment with compound **1** and daptomycin or vancomycin on clinically relevant MRSA strains. Heatmaps depict the outputs of the checkerboard assays used for evaluating antibacterial drug interaction. Each square represents quantitative data expressed as the mean value of optical density (OD_600 nm_, n = 3). Dark blue regions and white regions represent highest and lowest bacterial growth densities, respectively. (A-B) clinical isolate D15 (CA-MRSA); (C-D) clinical isolate D17 (HA-MRSA); (E-F) reference MRSA strain USA300. The FIC index (FICI) was calculated as the sum of the individual FICs of the antibacterial compounds under test in each well. FICI values were interpreted as follows: synergistic (Syn = FICI ≤ 0.5); additive (Add = 0.5 < FICI ≤1); indifferent (Ind = 1 < FICI ≤ 4); or antagonistic effect (Ant = FICI >4).

Lastly, we performed a preliminary toxicological assessment of compound **1** on human red blood cells (hRBCs) (Figure S15). The concentration causing 50% haemolysis (HEC□□) was 34.54 μg/mL, while daptomycin and vancomycin showed no hemolysis up to 200 μg/mL. Notably, the HEC□□ of compound **1** is about four times higher than its MBC against MRSA and MSSA strains, indicating moderate hemolysis (∼23%) at the bactericidal concentration of 8 μg/mL. Compared to other antibacterial secondary metabolites, the haemolytic activity of compound **1** at this concentration is considered moderate^51^.

## 4. Discussion

Decalin-containing tetramic acids (DTAs), such as compound **1**, have attracted widespread attention due to their potent antibacterial activity against various bacterial strains, such as *B. subtilis*, MRSA, vancomycin-resistant *Enterococcus*, and multidrug-resistant *Mycobacterium tuberculosis*^46,52,53^. Recent research has explored the structure-activity relationship of the antibacterial activity of DTA derivatives against MRSA, identifying key chemical features associated with antimicrobial potency^46^. For instance, dehydrogenation of the tyrosine sidechain of the tetramic acid substructure in conipyridoins, and the configuration at the C5’ position in 5’-epi-equisetin are crucial for antimicrobial activity against MRSA^46,48^. For several DTAs, such as N-demethylophiosetin, paecilosetin, and 8-epi-equisetin, the absence of oxygen groups as well as the stereochemistry of the methyl groups at the C8 position on the decalin skeleton are important determinants of the antibacterial activity of DTAs^46,48^. In cordysetin A, the D configuration of serine and N-methylation of the tetramic acid are crucial for its anti-tuberculosis activity^53^. Conversely, the antifungal activity of fusaramin appears to be more dependent on the identity of the amino acid than its cyclization to tetramic acid^54^.

DTAs with planar structures similar to compound **1**, such as CJ-21058 and the hyalodendrins A and B, have been relatively underexplored, with most studies focusing primarily on their chemical structures and larvicidal activities, while their antibacterial properties remain largely uninvestigated^41,42^. Therefore, our in-depth bioactivity investigation of compound **1** provides interesting new information. Our observation that it exerts strong antibacterial effects on Gram-positive bacteria, while showing no inhibitory effects against Gram-negative bacteria is consistent with previously reported DTAs^46^. Compared to the MIC values of other DTAs against MRSA (0.002-42 µg/mL)^46^, compound **1** also shows excellent antibacterial effects, with MIC values of 0.5-1 µg/mL. In comparison to equisetin (MIC = 1.25 µg/mL)^52^, the presence or absence of the methyl group at the C4 position, as well as differences in stereo-centres, do not appear to significantly affect the antibacterial activity against the tested MRSA strains.

Our findings further show that compound **1** has significant bactericidal effect on *S. aureus*, as exemplified for the three MRSA strains USA300, D15 and D17, and the MSSA strain ATCC 29213. Notably, compound **1** exerts stronger bactericidal activity against these strains compared to the positive control antibiotics daptomycin and vancomycin at 8 µg/mL, and it reveals additive or indifferent effects in combination treatment with daptomycin or vancomycin against the three tested MRSA strains. The structurally related compound equisetin has been demonstrated to effectively eradicate MRSA or VRE with low-level antibiotic resistance and to synergize with colistin, although it also has an indifferent effect with other antibiotics, including daptomycin^55^. Furthermore, equisetin has been shown to efficiently eliminate intracellular *S. aureus* by enhancing the host autophagy and inducing mitochondria-mediated reactive oxygen species production to clear the infection, demonstrating remarkable anti-infective activity in a peritonitis mouse infection model^56^. Although the bactericidal mechanism of compound **1** requires further investigation, these findings suggest its potential as a valuable antibacterial agent. In addition to their antimicrobial activity, DTAs have been shown to possess a variety of other biological activities, for instance, antifungal^54^, anticancer^52,57–59^, anti-HIV^60^, pesticidal activity^61^, anti-obesity^62^, and anti-atherosclerosis effects^63^. These diverse biological activities open avenues for further exploration of the pharmacological potential of compound **1**.

In conclusion, our findings imply that compound **1** is a promising antibacterial agent that may help to overcome antibiotic resistance. Altogether, this study not only presents a comprehensive genomic and metabolomic characterization of an underexplored fungus that has led to the identification of a potent antibacterial agent.

## Abbreviations

A domain: adenylation domain
Aas: auxiliary activities
ACP: acyl-carrier protein
AT: acyl-transferase
ATCC: American type culture collection
BGCs: biosynthetic gene clusters
BLAST: basic local alignment search tool
C domain: condensation domain
CAZyme: carbohydrate-active enzyme
CBMs: carbohydrate-binding modules
CD: circular dichroism
CEs: carbohydrate esterases
CFUs: colony-forming units
cMT: carbon-methyltransferase
DAse: diels-alderases
DH: dehydratase
DMSO: dimethyl sulfoxide
DPY: dextrose peptone yeast
DTAs: decalin-containing tetramic acids
ECD: electronic circular dichroism
EtOAc: ethyl acetate
FICI: fractional inhibitory concentration index
GenSAS: genome sequence annotation server
GHs: glycoside hydrolases
GNPS: global natural products social molecular networking
GO: gene ontology
GTs: glycosyl transferases
hRBCs: human red blood cells
LC-MS/MS: high-resolution, high-performance liquid chromatography-coupled tandem mass spectrometry
HR-PKS: highly-reducing polyketide synthase
ITS: internal transcribed spacer
KR: ketoreductase
KS: ketosynthase
LSU: large ribosomal subunit
MBC: minimum bactericidal concentration
MEA: malt extract agar
MICs: minimum inhibitory concentrations
ML: Maximum Likelihood
MRSA: methicillin-resistant *S. aureus*
MSSA: methicillin-sensitive *S. aureus*
NCBI: national center for biotechnology information
NCTC: national collection of type cultures
NMR: nuclear magnetic resonance spectroscopy
NR-PKS: non-reducing polyketide synthase
NRPS: non-ribosomal peptide synthetase
PCP: peptidyl carrier protein
PCR: polymerase chain reaction
PDA: potato dextrose agar
PKS: polyketide synthase
PLs: polysaccharide lyases
RiPP: post-translationally modified peptide
*Rpb2*: RNA polymerase II subunits 2
SAGM: Saline adenine glucose mannitol
SDA: Sabouraud dextrose agar
SMs: secondary metabolites
SNA: synthetically nutrient-poor agar
SSU: small ribosomal subunit
TD: terminal reductase
TE: thioesterase
*Tef1*: translation elongation factor 1α.

## Acknowledgments

The authors are grateful for technical assistance by Pieter Tepper, Rita Setroikromo, the staff at the interfaculty mass spectrometry center of the University Medical Center RUG/UMCG, bioinformatic support by Dr. Thomas Hackl (Groningen Institute for Evolutionary Life Sciences, RUG), NMR analysis support by Dr. Peter Fodran (Chemical and Pharmaceutical Biology, RUG), and ECD measurement support by Dr. Alexander Ryabchun (Faculty of Science and Engineering, Synthetic Organic Chemistry, RUG). We are grateful for Xiaofang Li in the Department of Medical Microbiology and Infection Prevention in UMCG for providing the MRSA strains.

## Supplementary Information

Supplementary Information file with Supplementary Methods and Results. Table S1. *Staphylococcus aureus* strains used for MBC tests in this study. Table S2. Reads and assembly statistics of *Neocucurbitaria*Lsp. VM-36. Table S3. Biosynthetic gene clusters of *Neocucurbitaria* sp. VM-36 predicted by antiSMASH version 7.0.1. Table S4. Compounds reported to be isolated from the Cucurbitariaceae family. Table S5. Molecular networking-based identification of secondary metabolites in *Neocucurbitaria* sp. VM-36. Table S6. MIC values of compound 1 against ESKAPE strains. Figure S1. Contigs of *Neocucurbitaria sp*. VM-36 visualized by Bandage. Figure S2. Functional genome annotation of *Neocucurbitaria* sp. VM-36. Figure S3. Structures of compound predicted to be produced by *Neocucurbitaria* sp. VM-36 based on antiSMASH analysis. Figure S4. Structures of compounds reported to be produced by fungi from the Cucurbitariaceae family. Figure S5. Total ion chromatograms recorded in positive ion mode of EtOAc extracts of *Neocucurbitaria* sp. VM-36 grown on PDA solid and DPY liquid medium. Figure S6. Calculated ECD spectra of the compound **1** isomer with C-1 being a carbonyl group and C-2’ being a hydroxyl group. Figure S7. ^1^H NMR spectrum of compound **1** in DMSO-d6 (600 MHz). Figure S8. ^13^C NMR spectrum of compound **1** in DMSO-d6 (150 MHz). Figure S9. HSQC spectrum of compound **1** in DMSO-d6. Figure S10. HMBC spectrum of compound **1** in DMSO-d6. Figure S11. ^1^H-^1^H COSY spectrum of compound **1** in DMSO-d6. Figure S12. NOESY spectrum of compound **1** in DMSO-d6. Figure S13. Amino acid sequence alignment between NeoC in *Neocucurbitaria* sp. VM-36 and Phm7 in *Pyrenochaetopsis* sp. RK10-F058. Figure S14. Effect of combination treatment with compound **1** and daptomycin or vancomycin on clinically relevant MRSA strains. Figure S15. *In vitro* hemolytic activity of compound **1** towards human red blood cells as compared to vancomycin and daptomycin.

## Data Availability statement

The sequencing data and genome assembly for this study have been deposited in the European Nucleotide Archive (ENA) at EMBL-EBI under the accession number PRJEB97715. The mass spectrometry data and the Cytoscape file of molecular networking have been deposited on GNPS under the accession number MassIVE ID: MSV000098902. The raw NMR data of compounds **1** is listed in the Supplementary Information. Antibacterial activity of compound **1**.

## Author Contributions

XL, TH, and KH designed the study and developed the workflow. XL and TH performed the genomic and metabolomic experiments. XL performed the purification and structural analysis of compound **1.** RdCFV and JMvD designed and RdCFV performed and analyzed the antibacterial assays and *in vitro* human toxicology assays. KH supervised the project. XL wrote the manuscript with input from all authors. All authors have read and approved the final version of this manuscript.

