## Supplementary Information for "Integrating genomics and metabolomics to accelerate the discovery of anti-MRSA natural products from the endophytic fungus *Neocucurbitaria* sp. VM-36"

[Evolutionary analysis of NeoC with some representative [4+2]-cyclases	3](#_Toc208992313)

### Supplementary Methods

#### Marfey’s method

To determine the configuration of amino acids in compound **1**, Marfey’s method was used according to the reported method with minor modification^1^. Compound **1** (0.2 mg) was dissolved in 2 mL of 6 M HCl and left at 110 ℃ for 16 h in a sealed tube. The hydrolysate was dried under a N_2_ flow. In an Eppendorf tube, 3.6 μmol of a 1% acetone solution of FDAA (N-(5-fluoro-2,4-dinitrophenyl)-l-alaninamide) and 20 μmol of a 1 M solution of NaHCO_3_ were added to the hydrolysate, and 2.5 μmol of standards D-serine and L-serine, respectively. The reaction mixture was heated with frequent shaking over a hot plate at 40 °C for 1 h and then cooled to RT. Then, 20 μmol of 2 M HCl and 1 mL of MeOH were added to the reaction mixture. The samples were analyzed by low-resolution LC-MS, and molecular weights and retention times were compared with those of standard serines. Acetonitrile/water containing 0.1% formic acid was used as the mobile phase under a linear gradient elution mode at a 0.5 mL/min flow rate. A mass range of *m/z* 100-1250 was covered with a scan time of 1 s, and data were collected in the positive ion mode. UV detection at 200-400 nm was performed by photodiode array detection.

#### Evolutionary analysis of NeoC with some representative [4+2]-cyclases

Sequences of multiple [4+2]-cyclases, including Phm7 from *Pyrenochaetopsis* sp. RK10-F058^2^, GNR600 from *F. heterosporum*^3^, PvhB from *Penicillium variabile*^4^, Fsa2 from *Fusarium* sp. FN080326^5^, Eqx3 from *F. heterosporum*^6^, CghA from *Chaetomium globosum*^7^, UcsH from *Acremonium* sp. KY4917^8^, MycB from *Myceliophthora thermophile*^9^, PoxQ from *P. oxalicum*^10^, PyiF from *Magnaporthe grisea* NI980^11^, CcsF from *Aspergillus clavatus* NRRL 1^12^, Pyri4 and Pyre3 from *Streptomyces rugosporus*^13^, were obtained from the UniProt or NCBI databases. They were aligned using ClustalW, and the Maximum Likelihood (ML) tree were generated in MEGA (v11) with 1000 bootstrap replicates. Enzymes marked with stars in Figure 10 of the main manuscript have reported crystal structures, and the remaining protein structures were predicted with AlphaFold3.

#### Antibacterial Susceptibility Assays

##### 1. Minimum Inhibitory Concentrations (MICs) and Minimum Bactericidal Concentrations (MBCs) of compound **1**

The broth microdilution method was used to determine the MIC values of compound **1** against different bacterial strains according to the guidelines from the European Committee for Antibacterial Susceptibility Testing (EUCAST, 2022) and previously reported references^14^. In brief, compound **1** was solubilized in DMSO, filtered with a 0.22 µm PTFE membrane, and serially diluted in sterile double-distilled water to obtain a concentration range of 0.03 - 64 µg/mL. Bacteria were reactivated on cation adjusted Mueller Hinton (MH) agar (Bioxon®) plates for 24 h at 36 ± 1º C. Overnight cultures were then prepared in cation adjusted MH broth at 36 ± 1 ºC with constant shaking at 250 revolutions per min (rpm), and added to 96-well microtiter plates at a final concentration of 5 $\times$ 10^4^ colony-forming units (CFUs)/well. Internal controls were used for each assay and included medium containing 10% DMSO (solvent control), medium with inoculum bacterial cells (bacterial growth control), and medium only (medium control) ^14^. The final concentration of DMSO in the assay was ≤ 0.6% *v*/*v*_final_. Plates were incubated in a Biotek® Epoch-2 microplate reader at 36 ± 1 ºC for 24 h with double-orbital agitation at a frequency of 282 cycles/min. The values of the lowest concentration of compound **1** that completely inhibited bacterial growth were determined as the MICs ^14^. The MIC values of ESKAPE strains were firstly evaluated (Table S5).

Due to the strong antibacterial effect of compound **1** on MRSA strains, we further determined the MBC values of this compound against 4 MRSA strains, namely *S. aureus* USA300, D15 and D17, and NE1688 (Sle1). The *S. aureus* ATCC 29213 strain was also included as a MSSA reference. The glycopeptide vancomycin and the cyclic lipopeptide daptomycin were used as positive controls. A volume of 2.5 µL of bacterial broth was taken from the MIC assay wells, with corresponding concentrations of 4 $\times$MIC, 2 $\times$MIC, and MIC, and then used to inoculate MH agar plates for MBC determination upon incubation at 36 ± 1 ºC for 24 h. Bacteria treated with the respective vehicles for compound **1**, vancomycin, and daptomycin served as negative controls (DMSO ≤0.6% v/v_final,_ double-distilled water, and NaCl 0.9% w/v, respectively). The MBC was determined as the lowest concentration of compound **1** that reduced the viable bacterial count on the agar plates by 99.9%^14^. All tests were performed in triplicates.

##### 2. Bacterial growth assay after compound 1 treatment

To monitor the effects of compound **1** on different MRSA and MSSA strains growth experiments were performed, using vancomycin and daptomycin as controls. In brief, the strains were reactivated, cultured, and seeded in 96-well microtiter plates as described above. After 10 h incubation, bacteria were treated with compound **1** or the control antibiotics at final concentrations of 2, 4, and 8 µg/mL, and incubated at 36 ± 1 ºC. Bacteria treated with the respective vehicles for compound **1**, vancomycin, and daptomycin served as negative controls (DMSO ≤0.6% v/v_final,_ double-distilled water, and NaCl 0.9% w/v, respectively). All tests were performed in triplicate (n = 3). The bacterial growth was monitored by OD_600_ measurements at 20 min intervals and results are presented using Graphpad^®^. The results are expressed as follows:

Bacterial growth (OD_600_) = (mean OD_600_ value compound-treated or untreated bacteria - mean OD_600_ value of vehicle treated bacteria).

The viability of bacteria was measured before (t = 8 h) and after 40 h compound exposure (t = 48 h). Briefly, 2 µL of bacterial broth was taken from each well, diluted (1: 10^6^ dilution factor), and then 2.5 µL of the dilutions were seeded onto MH agar plates, which were incubated at 36 ± 1 ºC for 24 h before CFUs were counted. The amount of CFUs/mL was calculated.

##### 3. Fractional inhibitory concentration index (FICI) Determination

We also evaluated the drug interaction of compound **1** (**A**) with daptomycin (**B**) or vancomycin (**C**) using a checkerboard assay in 96-well microtiter plates against three strains, including *S. aureus* USA300, D15 and D17. In total, 22 combinations were tested, which included five concentrations of compound **1** (final concentrations 2-0.13 µg/mL) together with seven concentrations of daptomycin or vancomycin (final concentrations 2-0.03 µg/mL). All treatments were tested in 3 independent experiments (n = 3). The assay was performed at a final concentration of 5 $\times$10^4^ CFUs/ well for each strain. After 24 h of incubation at 36 ± 1 ºC, MIC values of drug **A**, drug **B**, drug **C**, and their combinations were determined. FICI was calculated based on previously reported methods^15,16^, where FIC**A** = MIC of **A** in a combination with **B** /MIC of compound **A**, FIC**B** = MIC of **B** in a combination with **A** /MIC of compound **B**, and FICI = FIC**A** + FIC**B**. The same calculation was used for the combinations of **A** and **C**. These values were analysed by nonparametric models based on the Loewe additivity model (LA)^17^. FICI results were interpreted as synergistic (FICI ≤ 0.5), additive (0.5 < FICI ≤ 1), indifferent (1 < FICI ≤ 4) or antagonistic (FICI > 4)^18,19^. The lowest values of FICI (FICI_Min_), the median FICI (FICI_Med_), and the highest FICI (FICI_Max_) were calculated for each combination and in each strain. Outcomes of these experiments were plotted in heatmaps in Figure S13 using GraphPad Prism version 8.0.1.

##### 4. Hemolysis assay with human erythrocytes

Human red blood cells (hRBCs) were isolated from approximately 10 mL of whole blood from healthy donors using Lymphoprep buffer^TM^ (Stem cell technologies, Canada) following the manufacturers protocol with some modifications in the centrifugation parameters (1500 rpm, 30 min, 19 ℃). The fraction corresponding to hRBCs and neutrophils (aprox. 4 mL) was washed with 1 mL of 150 mM NaCl by centrifugation at 500$\times$ g, 5 min and 20 °C. Then, stock solutions of hRBCs at 2% (v/v) hRBCs were preserved at 4 °C for a maximum of 4 weeks. These were made by mixing 1 mL of the fraction from the previous stem with 49 mL of modified ‘saline adenine glucose mannitol’ (SAGM, final pH 6.1) medium^20^ prepared in Dulbecco’s phosphate-buffered saline (DPBS) buffer. hRBCs were allowed to precipitate in the storage media for 48 h before using them for bioassays.

On the day of experiments, the hRBCs stock was washed three times with DPBS (pH 7.2) by centrifuging (500$\times$ g, 5 min, 20 °C) to remove the SAGM medium. Subsequently, the hRBCs were resuspended in DPBS. 180 µL of a 1% (v/v) solution of intact hRBCs in DPBS (pH 7.2) were loaded into the wells of a U-shaped, 96-well microplate Greiner®. Then the hRBCs were treated with four concentrations of compound **1** (final in-test concentrations 200, 100, 10, 1 µg/mL). Also, four concentrations of the compound’s vehicle (DMSO at in-test concentrations 1%, 0.5% 0.05% and 0.005% v/v) were evaluated to be used as blanks. Treatment of hRBCs with a solution of sodium dodecyl sulfate (SDS) at a final in-test concentration 0.1% w/v was used as positive control for hemolysis, whereas addition of DPBS pH 7.2 served as negative control. Vancomycin and daptomycin were included in the evaluation for comparative purposes at the same final in tests concentrations as used for compound **1**. Their vehicles, double-distilled water or NaCl 0.9% w/v, respectively, were also evaluated as their blanks. Afterwards, the plates containing treated hRBCs were incubated at 37 °C for 24 h, and the microplate was then centrifuged at 500 rpm (10 min, 26 °C). Subsequently, 50 µL of the supernatant from the treated hRBCs were transferred into a new flat-bottomed 96-well plate containing 150 µL of DPBS pH 7.2. To quantify hemolysis, the plate was read at an optical density of 410 nm (OD_410 nm_) in a Biotek® instrument. The hemolytic activity was described in terms of percentage of hemolysis (% of hemolysis) and was calculated considering the Standard Practice for Assessment of Hemolytic Properties of Materials (ASTM F-756-00), using a previously reported equation^21^ adapted to our assay conditions as: 100 $\times$ (OD_410nm_ compound-treated hRBCs - OD_410nm_ vehicle-treated hRBCs)/ (OD_410nm_ of positive control - OD_410nm_ negative control). The results were then reported as the arithmetic mean of the percentage of hemolysis ± standard deviation obtained from each treatment. Additionally, the concentration of the antibacterial compound capable of inducing a hemolytic effect in 50% of the hRBCs (HEC_50_) was obtained through linear regression analysis from 4-point regression plots obtained in Graphpad Prism® v. 8.0.1. Each treatment was tested in three technical replicates.

### Supplementary Results

#### Note S1. Comparative Analysis and Biosynthetic Gene Cluster (BGC) Prediction

**Choline, metachelin, and scytalone/T3HN.** BGCs of *Neocucurbitaria* sp. VM-36 with choline (Region 3.3), metachelin C (Region 7.1), and scytalone/T3HN (Region 7.4) annotations are likely closely related to fungal development.

Choline is suspected to play a vital role in the growth of filamentous fungi and the regulation of mycelial morphology^22^. The cluster in region 3.3 of the *Neocucurbitaria* sp. VM-36 genome shows 100% identity with choline BGC from *Aspergillus nidulans* FGSC A4.

Metachelin C on the other hand is a coprogen siderophore, which is a small molecular iron chelator that participates in multiple cellular processes in fungi^23^. In *Metarhizium robertsii ARSEF* 23 growth in iron-deficient media was shown to lead to significantly up-regulated expression of two highly homologous genes *mrsidD* and *mrsidA* (NRPS core gene) and production of a series of coprogens and dimerumic acids, including metachelin C, metachelin A, metachelin A-CE, metachelin B, dimerumic acid 11-mannoside, and dimerumic acid^23^. These responses were proposed to relate to the maintenance of normal physiology and adaptation to adverse environments^23^. In the 7 *Cucurbitariaceae* fungi that we analyzed in this study, four genes, encoding an NRPS, an ABC transporter-related protein, a putative siderophore biosynthesis protein, and an AMP-dependent synthetase and ligase, are conserved and show high similarity to the biosynthetic genes for metachelin C from *Metarhizium robertsii ARSEF* 23^23^. In the *Neocucurbitaria* sp. VM-36 genome, these genes are located in region 7.1.

Lastly, scytalone/T3HN are precursors for the biosynthesis of the fungal 1,8-dihydroxynaphthalene melanin pigment that plays important roles in fungal development, survival, colorization, UV protection, oxidative stress and pathogenesis, as well as virulence in plant and human hosts^24–26^. The cluster in region 7.4 of the *Neocucurbitaria* sp. VM-36 genome exhibits 40% identity with the scytalone/T3HN biosynthetic gene cluster from *Pestalotiopsis fici W106-1* and is therefore likely to produce these compounds. Notably, in our morphological characterization of *Neocucurbitaria* sp. VM-36, we observed that the fungus produces chlamydospores and yellow pigment upon 28 days of culture in SDA medium. Moreover, upon incubation for 77 days, black pigment was produced in high amounts, which was accompanied by the growth of hyphae, the production of conidia, and an increase in colony size. Thus, it is possible that the scytalone/T3HN cluster in *Neocucurbitaria* sp. VM-36 is involved in helping the fungus to resist biotic or abiotic stresses and is responsible for significant morphological changes. However, these assumptions require further exploration.

**Xanthones.** Natural products with a xanthone scaffold are widely distributed in higher plants, lichens, bacteria, and fungi^27,28^. Multiple substitution positions and the ability to multimerize render this class of compounds highly diverse and complex. Accordingly, it has attracted attention due to a broad array of bioactivities, including cytotoxic, antioxidant, antimicrobial, anti-fungal, anti-inflammatory, antithrombotic activities^27–29^.

In the *Neocucurbitaria* sp. VM-36 genome, the cluster in region 13.3 shows similarity to a series of xanthone-related BGCs in the MIBiG database, including BGC0001886 (secalonic acids, 37% of genes show similarity), BGC0002063 (cryptosporioptide B, 23% of genes show similarity), BGC0001988 (neosartorin, 31% of genes show similarity), BGC0002726 (4-chloropinselin, 20% of genes show similarity), BGC0001403 (trypacidin, 28% of genes show similarity), BGC0000121 (RES-1214-2, 25% of genes show similarity), BGC0002592 (geodin, 33% of genes show similarity), BGC0002257 (rufoschweinitzin, 42% of genes show similarity), BGC0002244 (3’-methoxy-1,2-dehydropenicillide, 15% of genes show similarity), and BGC0002062 (agnestin A, 28% of genes show similarity). A common feature of these clusters is that the key non-reducing polyketide synthase (nr-PKS) can use acetyl and malonyl CoA to form C16 polyketides^29^ and it does not carry a thioesterase (TE) domain. Therefore, an additional metallo-β-lactamase-type thioesterase is needed for releasing the polyketide from the PKS^30^. With the function of other enzymes encoded in the clusters, the linear polyketides are subsequently converted into anthroquinone, benzophenone, and finally the xanthone skeleton^29^. These core enzymes are also encoded in the BGC in region 13.3 and share high amino acid sequence similarity with their homologues from literature mentioned above. In addition, genes annotated as short-chain dehydrogenase, FAD-binding domain, drug resistance transporter, NAD(P)H-binding, and oxidoreductase activity, also show high similarities to genes in multiple BGCs. Other biosynthetic genes in the BGC of region 13.3, however, are different from known xanthone-related BGCs, indicating that this BGC might produce previously unknown xanthone compounds. These ideas could be further investigated by heterologous expression.

**Griseovulvin analogs.** The cluster in region 15.2 of *Neocucurbitaria* sp. VM-36 is a hybrid of a T1PKS and fungal-Ripp-like BGC. The core PKS gene in this BGC shows 63% identity with the polyketide synthase in BGC0000070 of *Penicillium aethiopicum*. This BGC was previously shown to be responsible for production of the antifungal agents griseofulvin, epidechlorogriseofulvin, norlichexanthone, dehydrogriseofulvin, 4-desmethyl griseofulvin, and griseoxanthone B^31,32^. The PKS enzyme, GsfA, was demonstrated to use one acetyl-CoA and six malonyl-CoA to form a benzophenone precursor^32^. This precursor is converted into a spirocyclic or grisan scaffold by the cytochrome P450 GsfF, which performs the oxidative coupling between orcinol and phloroglucinol rings^32^. The cluster in *Neocucurbitaria* sp. VM-36 only shows similarity with the core gene and drug resistance transporter gene in the BGC from *P. aethiopicum*. The absence of additional enzymes and the presence of a fungal-Ripp-like gene suggests that this BGC has the potential to produce other complex compound skeletons^32,33^.

**Benzenediol lactones.** The cluster in region 4.2 of *Neocucurbitaria* sp. VM-36 shows a high degree of similarity to 7 known BGCs, including BGC0000076 (hypothemycin, 66% of genes show similarity), BGC0001245 (lasiodiplodin, 66% of genes show similarity), BGC0002187 (monorden D, 40% of genes show similarity), BGC0000134 (radicicol, 40% of genes show similarity), BGC0001246 (trans-resorcylide, 33% of genes show similarity), BGC0000045 (dehydrocurvularin, 25% of genes show similarity), and BGC0001057 (zearalenone, 18% of genes show similarity). The compounds produced from these BGCs are members of the benzenediol lactone family, a type of fungal polyketide metabolite possessing a macrolide core structure fused into a resorcinol aromatic ring. These compounds have been reported to possess several interesting biological activities, such as cytotoxicity, nematicidal properties, inhibition of various kinases, receptor agonists, anti-inflammatory activities, heat shock responses and immune system modulatory activities^34^.

The core gene in region 4.2 encodes a protein with highly-reducing (HR) PKS, glutathione S-transferase, and non-reducing PKS enzyme activities, sharing 63% identity with the protein AHV78247.1 in the MIBiG database, as well as 79% identity with AHV78243.1, and 70% identity with AHV78245.1 from *Lasiodiplodia theobromae*, respectively. Besides, genes with predicted cytochrome P450, EmrB/QacA drug resistance transporter, and O-methyltransferase functions also display high similarities with corresponding genes in the hypothemycin BGC of ACD39751.1, ACD39756.1, and ACD39755.1, respectively. Additional tailoring genes in the BGC of region 4.2 are likely to contribute to the final structures of the compounds produced.

In conclusion, although some BGCs have been annotated, 23 of the 34 BGCs of *Neocucurbitaria* sp. VM-36 remain unknown. They are potential treasure troves for drug discovery and need to be further explored.

#### Note S2. Molecular Networking-Based Secondary Metabolite Identification

The major constituents with high abundance in the crude extracts of *Neocucurbitaria* sp. VM-36 were annotated to be CJ-21058 analogues in cluster 5 (Figure 5B), which are tetramic acid type of components containing decalin rings. Different amino acids participating in the formation of the tetramic acid moiety and different side chain groups in the decalin rings enrich the diversity of this type of compounds. The highly abundant metabolite *m/z* 388.248 is annotated as [M + H]^+^ adduct of CJ-21058 (**1**) and *m/z* 387.247 is the isotope peak, with a mass difference of -1.001 Da. Although nodes *m/z* 387.993 and *m/z* 388.487 display different precursor mass, their MS2 fragments are the same as those in *m/z* 387.247, indicating that these nodes are CJ-21058 isomers. They all are highly abundant in crude extracts of *Neocucurbitaria* sp. VM-36. Here, it is worth noting that stereoselective structural diversity has been previously reported in this type of compounds ^35^. CJ-21058 has six stereoselective positions, which could lead to the formation of isomers with the same planar structures but different configurations. CJ-21058 was firstly reported in 2002 without determined stereoselective positions^36^. Interestingly, two new decalin/tetramic acid hybrid fungal metabolites hyalodendrin A (2*R*, 3*S*, 6*R*, 8*S*, 11*S*, 5’*R*) and hyalodendrin B (2*R*, 3*S*, 6*R*, 8*S*, 11*S*, 5’*S*) were reported in 2020^36,37^. They show the same planar structure as CJ-21058, and possess the opposite configuration only at the chiral center CH-5’^36,37^. Therefore, it is highly likely that these high abundant nodes in cluster 5 have the same planar structures as CJ-21058, but with different stereoselectivities. In addition, other nodes are also predicted according to edge annotations in this cluster. Node **20** (*m/z* 402.264) has one more -CH_2_ group than **1**, and one less -C atom than node **21** (*m/z* 414.264). Node **22** (*m/z* 400.246) has one less -CH_2_ group than **21**, and it does not have a direct relationship with node **20**. According to their MS2 fragments in Table S4, **21** is predicted to be the [M + H]^+^ adduct of (E)-3-(hydroxy(1,3,6-trimethyl-2-((1E,3E)-penta-1,3-dien-1-yl)-1,2,4a,5,6,7,8,8a-octahydronaphthalen-1-yl)methylene)-5-(hydroxymethyl)-1-methylpyrrolidine-2,4-dione or LL-49F233alpha or phomasetin, which have the same planar structure. Node **20** has a side chain group in the decalin ring that differs from **21**, and it is predicted to be (E)-3-((2-((E)-but-1-en-1-yl)-1,3,6-trimethyl-1,2,4a,5,6,7,8,8a-octahydronaphthalen-1-yl)(hydroxy)methylene)-5-(hydroxymethyl)-1-methylpyrrolidine-2,4-dione, which is a putative new compound. Compared with **21**, node **22** lacks an N-methyl group in the tetramic acid moiety, and is identified as (E)-3-(hydroxy(1,3,6-trimethyl-2-((1E,3E)-penta-1,3-dien-1-yl)-1,2,4a,5,6,7,8,8a-octahydronaphthalen-1-yl)methylene)-5-(hydroxymethyl)pyrrolidine-2,4-dione or N-demethylated-LL-49F233alpha or N-demethylated-phomasetin. Both node **23** (*m/z* 374.233) and node **24** (*m/z* 374.232), with ions [M + H]^+^, show one less -CH_2_ group than **1**. According to the spectral fragments, the existence of fragments (*m/z* 217.1953 and *m/z* 245.1908) in the node **23** indicate that its side chains in the decalin ring are the same as those in node **1**, while the absence of fragment (*m/z* 126.0554) reveals that there is no -N-CH_3_ group in the tetramic acid part. Therefore, node **23** is predicted to be (E)-3-(hydroxy(1,3,6-trimethyl-2-((E)-prop-1-en-1-yl)-1,2,4a,5,6,7,8,8a-octahydronaphthalen-1-yl)methylene)-5-(hydroxymethyl)pyrrolidine-2,4-dione, the N-demethylated analogue of CJ-21058. On the contrary, node **24** does not have fragments (*m/z* 217.1953, *m/z* 245.1908, and *m/z* 328.2273). Instead, the presence of fragments (*m/z* 126.0554 and *m/z* 144.0657) indicates the absence of -CH_2_ in the decalin ring and the presence of a -N-CH_3_ group in the tetramic acid part. According to the presence of *m/z* 288.1593 and absence of *m/z* 304.1536, the -CH_3_ group attached to position 4 is lost and node **24** is annotated to be (E)-3-((1,6-dimethyl-2-((E)-prop-1-en-1-yl)-1,2,4a,5,6,7,8,8a-octahydronaphthalen-1-yl)(hydroxy)methylene)-5-(hydroxymethyl)-1-methylpyrrolidine-2,4-dione, which has a similar planar structure as equisetin. Compared with **1,** nodes **25** (*m/z* 358.238) and **26** (*m/z* 370.238) have lost -CH_2_O and -H_2_O groups, respectively. Node **25** has lost an -O atom and a -C atom compared with node **23** and **26**. The significantly increased fragments with *m/z* 342.2430 and *m/z* 152.0708 in node **26** and *m/z* 340.2264, *m/z* 330.2433 and *m/z* 140.0708 in node **25**, and the sharply reduced fragment *m/z* 170.0813 in both nodes indicate changes in the tetramic acid part, but not in the decalin part. Node **25** and **26** represent two putative new compounds, which are predicted to be [M + H]^+^ adducts of (Z)-3-(hydroxy(1,3,6-trimethyl-2-((E)-prop-1-en-1-yl)-1,4,4a,5,6,7,8,8a-octahydronaphthalen-1-yl)methylene)-1-methyl-1,3-dihydro-2H-4l5-pyrrole-2,4-dione and (E)-3-(hydroxy(1,3,6-trimethyl-2-((E)-prop-1-en-1-yl)-1,2,4a,5,6,7,8,8a-octahydronaphthalen-1-yl)methylene)-1-methyl-5-methylenepyrrolidine-2,4-dione, respectively.

### Supplementary Tables

#### Table S1. *Staphylococcus aureus* strains used for MBC tests in this study.

| **Bacterial strain** | **Genotype/description** | **Reference** |
| --- | --- | --- |
| *S. aureus* D15 (CA) | Community-associated (CA) clinical MRSA isolate. More information is available at <https://www.ebi.ac.uk/ena/browser/view/PRJEB17079> | ^38^ |
| *S. aureus* D17 (HA) | Hospital-associated (HA) MRSA isolate. More information is available at https://www.ebi.ac.uk/ena/browser/view/PRJEB17079 |  |
| *S. aureus* ATCC 29213 | /MSSA | - |
| *S. aureus* USA300 | USA300 wild-type, MRSA | ^39^ |
| *S. aureus* NE1688 (Sle1) | USA300 JE2 derivative with a transposon insertion in SAUSA300_0438, Em^R^ | ^40^ |

#### Table S2. Reads and assembly statistics of *Neocucurbitaria* sp. VM-36*.*

|  | **Unfiltered** | **Filtered (>2 kb)** |
| --- | --- | --- |
| No. of bases (Gb) | 5.5 | 5.2 |
| No. of reads | 870,296 | 590,512 |
| N50 (kb) | 5.5 | 7.3 |
| Mean read length (bp) | 6,422.9 | 9,074.5 |
| Mean read quality | 16.1 | 17.1 |
| Coverage | 130× | |
| Polishing steps | Medaka | |
| No. of contigs (coverage ≥ 40×) | 19 | |
| Genome size (Mb) | 32.8 | |
| GC (%) | 50 | |
| BUSCO (Ascomycota_odb10) (%) | 97.2 | |
| Number of the protein-coding genes | 11,817 | |
| tRNA genes | 120 | |
| rRNA genes | 29 | |
| Proteins with a predicted Pfam domain | 9,806 | |
| CAZymes | 570 | |

#### Table S3. Biosynthetic gene clusters of *Neocucurbitaria* sp. VM-36 predicted by antiSMASH version 7.0.1.

| **Cluster ID** | **Region** | **Contig** | **Type** | **From (nt)** | **To (nt)** | **Most similar known cluster** | **Genbank** | **Type** | **Similarity** |
| --- | --- | --- | --- | --- | --- | --- | --- | --- | --- |
| 1 | 2.1 | 21 | Fungal-Ripp-like | 61425 | 152320 |  |  |  |  |
| 2 | 2.2 | 21 | NRPS-like | 882440 | 944272 |  |  |  |  |
| 3 | 2.3 | 21 | NRPS | 1287667 | 1352639 |  |  |  |  |
| 4 | 3.1 | 18 | Fungal-Ripp-like | 1044648 | 1132776 |  |  |  |  |
| 5 | 3.2 | 18 | Fungal-Ripp-like | 1616536 | 1707578 |  |  |  |  |
| 6 | 3.3 | 18 | NRPS-like | 1949834 | 2008639 | Choline | CH236925.1 | NRP | 1 |
| 7 | 3.4 | 18 | NRPS | 2381577 | 2450639 |  |  |  |  |
| 8 | 4.1 | 24 | Fungal-Ripp-like | 236535 | 321230 |  |  |  |  |
| 9 | 4.2 | 24 | T1PKS | 2457213 | 2533431 | Hypothemycin | EU520417.1 | Polyketide | 0.66 |
| 10 | 4.3 | 24 | Fungal-Ripp-like | 3165551 | 3256546 |  |  |  |  |
| 11 | 5.1 | 39 | Terpene | 1212913 | 1239972 | Squalestatin S1 | LDZW01000177.1 | Terpene | 0.40 |
| 12 | 5.2 | 39 | Terpene | 1648619 | 1680518 |  |  |  |  |
| 13 | 7.1 | 31 | NRPS | 119867 | 183763 | Metachelin C/ metachelin A/ metachelin A-CE/ metachelin B/ dimerumic acid 11-mannoside/ dimerumic acid | ADNJ02000002.1 | NRP | 0.5 |
| 14 | 7.2 | 31 | Fungal-Ripp-like | 997021 | 1088115 |  |  |  |  |
| 15 | 7.3 | 31 | NRPS, T1PKS | 2001288 | 2073239 | Phyllostictine A/B | KY682688.1 | NRP + Polyketide | 0.2 |
| 16 | 7.4 | 31 | T1PKS | 2492633 | 2559242 | Scytalone/T3HN | KI912112.1 | Polyketide | 0.4 |
| 17 | 8.1 | 7 | Fungal-Ripp-like | 86664 | 177673 |  |  |  |  |
| 18 | 8.2 | 7 | NRP-metallophore, NRPS | 270853 | 356969 |  |  |  |  |
| 19 | 8.3 | 7 | T1PKS | 437365 | 505222 |  |  |  |  |
| 20 | 8.4 | 7 | Terpene | 724153 | 760793 |  |  |  |  |
| 21 | 10.1 | 17 | T1PKS | 525493 | 595142 | (-)-Mellein | KM365454.1 | Polyketide | 1 |
| 22 | 13.1 | 13 | NRPS, T1PKS | 97521 | 169851 | Phomasetin | LC361337.1 | NRP + Polyketide | 0.85 |
| 23 | 13.2 | 13 | Fungal-Ripp-like | 636873 | 727937 |  |  |  |  |
| 24 | 13.3 | 13 | T1PKS | 820674 | 886262 | Secalonic acids | CAGA01000032.1 | Polyketide | 0.37 |
| 25 | 15.1 | 11 | T1PKS | 257292 | 323502 |  |  |  |  |
| 26 | 15.2 | 11 | T1PKS, fungal-Ripp-like | 2976045 | 3120187 | Griseofulvin / Epidechlorogriseofulvin / Norlichexanthone / Dehydrogriseofulvin / 4-desmethylgriseofulvin/ Griseoxanthone B | GU574478.1 | Polyketide: Iterative type I | 0.09 |
| 27 | 17.1 | 19 | NRPS-like | 1170174 | 1233809 |  |  |  |  |
| 28 | 18.1 | 16 | NRPS-like | 238848 | 302100 |  |  |  |  |
| 29 | 20.1 | 28 | T1PKS | 62317 | 136949 |  |  |  |  |
| 30 | 20.2 | 28 | T1PKS | 1167883 | 1219333 | Usnic acid | MG777490.1 | Polyketide | 0.33 |
| 31 | 21.1 | 25 | NRPS | 629575 | 712040 |  |  |  |  |
| 32 | 21.2 | 25 | Fungal-Ripp-like | 1513620 | 1604292 |  |  |  |  |
| 33 | 21.3 | 25 | T1PKS | 1864153 | 1926374 |  |  |  |  |
| 34 | 21.4 | 25 | T1PKS | 3492386 | 3559957 |  |  |  |  |

#### Table S4. Compounds reported to be isolated from the Cucurbitariaceae family.

| **No.** | **Name** | **Chemical formula** | **Species** | **Ref.** |
| --- | --- | --- | --- | --- |
| 1 | 8a-hydroxy-spiciferinone | C_14_H_18_O_4_ | *Pyrenochaeta nobilis* | ^41^ |
| 2 | A32390A | C_18_H_24_N_2_O_8_ | *Pyrenochaeta sp.* | ^42^ |
| 3 | Pyrenochaetamide A | C_18_H_16_N_2_O_4_ | *Pyrenochaeta sp.* | ^43^ |
| 4 | Pyrenochaetic acid A | C_13_H_14_O_4_ | *Pyrenochaeta terrestris* | ^44^ |
| 5 | Pyrenochaetic acid B | C_13_H_16_O_5_ | *Pyrenochaeta terrestris* | ^44^ |
| 6 | Pyrenochaetic acid C | C_13_H_16_O_4_ | *Pyrenochaeta terrestris* | ^44^ |
| 7 | Pyrenochaetolide A | C_18_H_18_O_7_ | *Pyrenochaeta sp.* | ^43^ |
| 8 | Pyrenochaetolide B | C_10_H_14_O_4_ | *Pyrenochaeta sp.* | ^43^ |
| 9 | Pyrenochaetoxy A | C_18_H_20_O_6_ | *Pyrenochaeta sp.* | ^43^ |
| 10 | Pyrenocine C | C_11_H_14_O_4_ | *Pyrenochaeta terrestris* | ^43^ |
| 11 | Neocucurbol A | C_20_H_28_O_4_ | *Neocucurbitaria unguis-hominis* FS685 | ^45^ |
| 12 | Neocucurbol B | C_20_H_28_O_4_ | *Neocucurbitaria unguis-hominis* FS685 | ^45^ |
| 13 | Neocucurbol C | C_20_H_28_O_4_ | *Neocucurbitaria unguis-hominis* FS685 | ^45^ |
| 14 | Neocucurbol D | C_20_H_28_O_3_ | *Neocucurbitaria unguis-hominis* FS685 | ^45^ |
| 15 | Neocucurbol E | C_20_H_32_O_2_ | *Neocucurbitaria unguis-hominis* FS685 | ^45^ |
| 16 | Neocucurbol F | C_20_H_34_O_3_ | *Neocucurbitaria unguis-hominis* FS685 | ^45^ |
| 17 | Neocucurbol G | C_20_H_32_O_2_ | *Neocucurbitaria unguis-hominis* FS685 | ^45^ |
| 18 | Neocucurbol H | C_20_H_32_O_3_ | *Neocucurbitaria unguis-hominis* FS685 | ^45^ |
| 19 | Neocucurbin A | C_20_H_30_O_5_ | *Neocucurbitaria unguis-hominis* FS685 | ^46^ |
| 20 | Neocucurbin B | C_20_H_30_O_5_ | *Neocucurbitaria unguis-hominis* FS685 | ^46^ |
| 21 | Neocucurbin C | C_21_H_32_O_4_ | *Neocucurbitaria unguis-hominis* FS685 | ^46^ |
| 22 | Neocucurbin D | C_20_H_32_O_4_ | *Neocucurbitaria unguis-hominis* FS685 | ^46^ |
| 23 | Neocucurbin E | C_22_H_34_O_5_ | *Neocucurbitaria unguis-hominis* FS685 | ^46^ |
| 24 | Neocucurbin F | C_19_H_28_O_5_ | *Neocucurbitaria unguis-hominis* FS685 | ^46^ |
| 25 | Neocucurbin G | C_19_H_28_O_4_ | *Neocucurbitaria unguis-hominis* FS685 | ^46^ |
| 26 | Phomalevone A | C_30_H_26_O_10_ | *Phoma sp.* | ^47^ |
| 27 | Phomalevone B | C_30_H_26_O_10_ | *Phoma sp.* | ^47^ |
| 28 | Phomalevone C | C_30_H_24_O_10_ | *Phoma sp.* | ^47^ |

#### Table S5. Molecular networking-based putative identification of secondary metabolites in *Neocucurbitaria* sp. VM-36.

| **No.** | **Metabolite name** | **t_R_**  **(min)** | **Experi-mental mass (*m*/*z*)** | **Adduct** | **Molecular formula** | **Molecular weight** | **Exact mass (*m*/*z*)** | **MZ Error**  **PPM (ppm)** | **MS^2^ fragment ions (*m*/*z*)** | **Growth medium** | |
| --- | --- | --- | --- | --- | --- | --- | --- | --- | --- | --- | --- |
|  |  |  |  |  |  |  |  |  |  | **PDA** | **DPY** |
| 1 | CJ-21058 | 20.33 | 388.2493 | [M+H]^+^ | C_23_H_33_NO_4_ | 387.52 | 388.2478 | -3.86 | 370.2380, 360.2533, 346.2375, 340.2264, 328.1914, 316.1915, 314.1739, 304.1544, 302.1752, 300.1602, 292.1541, 290.1734, 288.1599, 280.1539, 276.1599, 274.1450, 262.1440, 252.1231, 250.1443, 245.1911, 219.1752, 217.1953, 198.0758, 191.1796, 189.1639, 182.0813, 170.0813, 163.1482, 161.1326, 156.0656, 147.1168, 144.0657, 140.0707, 137.1323, 135.1167, 126.0547, 121.1012, 109.1012, 95.0855, 81.0698 | + | + |
| 2 | FruLeuIle | 2.05 | 407.2399 | [M+H]^+^ | C_18_H_34_N_2_O_8_ | 406.48 | 407.2394 | -1.22 | 389.2281, 371.2182, 353.2076, 335.1962, 323.1968, 307.2026, 276.1463, 258.1340, 245.1859, 230.1388, 212.1283, 132.1020, 102.0554, 86.0963 | - | + |
| 3 | DG(16:1/18:2/0:0) | 23.22 | 608.5259 | [M+NH_4_]^+^ | C_37_H_66_O_5_ | 590.93 | 608.5245 | -2.30 | 573.4883, 337.2744, 311.2582, 237.2215, 137.1332, 121.1011, 111.1168, 97.1012, 95.0854 | - | + |
| 4 | DG(18:2/0:0/18:2) | 25.08 | 634.5416 | [M+NH_4_]^+^ | C_39_H_68_O_5_ | 616.97 | 634.5410 | -0.95 | 599.5033, 337.2740, 263.2373, 189.1641, 175.1481, 165.1280, 151.1122, 147.1169, 137.1326, 133.1013, 127.1116, 121.1013, 111.1168, 99.0803, 97.1012, 93.0699, 83.0855, 57.0702 | - | + |
| 5 | Asp-Leu | 1.62 | 247.1299 | [M+H]^+^ | C_10_H_18_N_2_O_5_ | 246.26 | 247.1288 | -4.45 | 201.1235, 181.0974, 155.1180, 132.1020, 116.0342, 102.0550, 88.0391, 86.0963, 74.0235, 70.0288 | - | + |
| 6 | AEG(o-16:3/15:0) | 24.61 | 535.4732 | [M+H]^+^ | C_34_H_62_O_4_ | 534.87 | 535.4725 | -1.31 | 257.2459, 149.1327, 147.1171, 137.1332, 135.1168, 123.1169, 121.1015, 109.1013, 97.1011, 95.0855, 83.0853, 81.0699, 71.0856, 69.0699, 67.0544, 57.0700 | - | + |
| 7 | 4-[5-[[4-[5-[acetyl(hydroxy)amino]pentylamino]-4-oxobutanoyl]-hydroxyamino]pentylamino]-4-oxobutanoic acid | 18.83 | 478.2882 | [M+NH_4_]^+^ | C_20_H_36_N_4_O_8_ | 460.53 | 478.2927 | 9.40 | 337.2738, 306.2797, 263.2365, 245.2262, 216.0642, 163.1483, 149.1327, 147.1164, 137.1324, 135.1171, 121.1014, 109.1012, 95.0855, 83.0853, 81.0698 | + | + |
| 8 | AEG(o-18:4/18:2) | 25.12 | 599.5045 | [M+H]^+^ | C_39_H_66_O_4_ | 598.95 | 599.5040 | -0.83 | 543.4783, 525.4657, 473.4034, 459.3823, 361.2741, 337.2747, 333.2796, 321.2787, 319.2646, 309.2791, 263.2371, 245.2276, 221.1899, 203.1799, 189.1636, 177.1636, 175.1484, 165.1642, 163.1482, 161.1328, 153.1275, 151.1482, 149.1327, 137.1326, 135.1169, 123.1169, 121.1013, 113.0960, 111.1170, 109.1012, 99.0805, 97.1011, 95.0855, 85.0646, 83.0855, 81.0698, 69.0699, 67.0543, 57.0701, 55.0545 | - | + |
| 9 | Riboflavin | 5.79 | 377.1467 | [M+H]^+^ | C_17_H_20_N_4_O_6_ | 376.37 | 377.1454 | -3.44 | 359.1352, 341.1257, 243.0878, 200.0821, 135.0648, 117.0545, 99.0440, 75.0440, 73.0283, 69.0335, 61.0286, 57.0338 | + | + |
| 10 | Isousnic acid | 16.70 | 377.1242 | [M+H+CH_3_OH]^+^ | C_18_H_16_O_7_ | 344.32 | 377.1242 | 0 | 345.0982, 327.0874, 317.1031, 303.0871, 285.0754, 275.0919, 261.0769, 251.0917, 235.0982, 233.0816, 219.0662, 85.0285 | + | - |
| 11 | N-[3-[5,8-bis[3-[acetyl(hydroxy)amino]propyl]-14-(hydroxymethyl)-3,6,9,12,15,18-hexaoxo-1,4,7,10,13,16-hexazacyclooctadec-2-yl]propyl]-N-hydroxyacetamide/ desferriferricrocin | 2.80 | 718.3377 | [M+H]^+^ | C_28_H_47_N_9_O_13_ | 717.73 | 718.3379 | 0.27 | 546.2548, 528.2414, 471.2210, 402.1983, 374.1669, 356.1574, 345.1785, 317.1454, 314.1433, 303.1650, 299.1366, 285.1560, 275.1345, 272.1255, 230.1141, 190.1179, 188.1032, 173.0922, 145.0973, 131.0815, 114.0549, 86.0600, 70.0650 | - | + |
| 12 | Linoleic acid | 21.10 | 281.2486 | [M+H]^+^ | C_18_H_32_O_2_ | 280.45 | 281.2477 | -3.20 | 263.2372, 221.2270, 195.1591, 193.1591, 181.1591, 179.1433, 169.1221, 167.1435, 165.1275, 153.1277, 139.1119, 129.0909, 125.1327, 115.0753, 111.1169, 97.1012, 85.1012, 57.0702 | + | + |
| 13 | Arg-C18:2 | 15.45 | 437.3497 | [M+H]^+^ | C_24_H_44_N_4_O_3_ | 436.64 | 437.3488 | -2.05 | 420.3224, 419.3378, 378.3013, 239.1433, 175.1194, 158.0927, 116.0707, 115.0867, 114.1026, 112.0870, 70.0651 | - | + |
| 14 | 9,10-epoxy-12(Z)-octadecenoic acid | 16.28 | 279.2330 | [M+H-H_2_O]^+^ | C_18_H_32_O_3_ | 296.45 | 279.2319 | -3.93 | 261.2215, 209.1542, 195.1380, 181.1589, 167.1435, 165.1276, 163.1121, 153.1276, 151.1483, 137.1326, 123.1169, 113.0961, 99.0805, 97.1012, 85.1012, 71.0855, 57.0702 | - | + |
| 15 | Gly-C18:2 | 20.32 | 338.2701 | [M+H]^+^ | C_20_H_35_NO_3_ | 337.50 | 338.2676 | -7.39 | 263.2377, 245.2268, 175.1481, 161.1326, 147.1169, 133.1014, 121.1012, 109.1012, 95.0855, 81.0698 | - | + |
| 16 | 9S-Hydroxy-10E,12Z,15Z-octadecatrienoic acid | 17.10 | 277.2173 | [M+H-H_2_O]^+^ | C_18_H_30_O_3_ | 294.44 | 277.2165 | -2.88 | 277.2164, 259.2060, 235.1698, 231.2105, 221.1531, 217.1953, 207.1383, 189.1665, 189.1276, 181.1220, 175.1485, 153.0910, 149.1328, 145.1012, 135.1170, 123.1170, 109.1012, 95.0855, 93.0699, 69.0699, 67.0543 | + | + |
| 17 | 1,4a-dimethyl-9-oxo-7-propan-2-yl-3,4,10,10a-tetrahydro-2H-phenanthrene-1-carboxylic acid | 17.36 | 315.1966 | [M+H]^+^ | C_20_H_26_O_3_ | 314.43 | 315.1955 | -3.48 | 269.1908, 227.1801, 199.1120, 187.1120, 147.0806, 69.0701 | - | + |
| 18 | Phytosphingosine | 14.71 | 318.3014 | [M+H]^+^ | C_18_H_39_NO_3_ | 317.51 | 318.3002 | -3.77 | 300.2899, 282.2793, 270.2793, 265.2532, 264.2687, 252.2688, 90.0549, 74.0600, 60.0446 | - | + |
| 19 | 1-Linoleoylglycerol | 20.71 | 355.2854 | [M+H]^+^ | C_21_H_38_O_4_ | 354.53 | 355.2847 | -1.97 | 337.2740, 263.2370, 247.2421, 179.1794, 165.1640, 153.1282, 151.1481, 141.1275, 137.1328, 125.1332, 113.0961, 111.1167, 97.1013, 93.0697, 85.0649, 71.0857, 57.0701 | - | + |
| **Putative structures according to molecular network** | | | | | | | | | | |  |
| 20* | (E)-3-((2-((E)-but-1-en-1-yl)-1,3,6-trimethyl-1,2,4a,5,6,7,8,8a-octahydronaphthalen-1-yl)(hydroxy)methylene)-5-(hydroxymethyl)-1-methylpyrrolidine-2,4-dione | 20.74 | 402.2650 | [M+H]^+^ | C_24_H_35_NO_4_ | 401.55 | 402.2643 | -1.74 | 384.2536, 374.2690, 372.2547, 328.1920, 318.1698, 314.1754, 304.1541, 292.1546, 290.1753, 288.1606, 266.1389, 259.2064, 231.2108, 217.1954, 203.1797, 189.1639, 177.1642, 170.0814, 161.1325, 144.0656, 135.1170, 109.1012, 100.0392, 95.0855, 83.0853 | + | + |
| 21* | (E)-3-(hydroxy(1,3,6-trimethyl-2-((1E,3E)-penta-1,3-dien-1-yl)-1,2,4a,5,6,7,8,8a-octahydronaphthalen-1-yl)methylene)-5-(hydroxymethyl)-1-methylpyrrolidine-2,4-dione/ LL-49F233alpha/ phomasetin | 20.96 | 414.2650 | [M+H]^+^ | C_25_H_35_NO_4_ | 413.56 | 414.2643 | -1.68 | 396.2534, 386.2700, 354.2072, 346.2014, 328.1902, 318.1700, 304.1549, 300.1590, 288.1595, 286.1424, 278.1393, 271.2066, 266.1383, 243.2108, 229.1949, 224.1281, 215.1797, 198.0791, 188.0556, 175.1483, 173.1331, 170.0448, 156.0656, 147.1167, 144.0657, 140.0339, 137.1325, 121.1013, 109.1014, 107.0856, 95.0853, 81.0699, 69.0700 | + | + |
| 22* | (E)-3-(hydroxy(1,3,6-trimethyl-2-((1E,3E)-penta-1,3-dien-1-yl)-1,2,4a,5,6,7,8,8a-octahydronaphthalen-1-yl)methylene)-5-(hydroxymethyl)pyrrolidine-2,4-dione/ N-demethylated-LL-49F233alpha/ N-demethylated- phomasetin | 20.52 | 400.2487 | [M+H]^+^ | C_24_H_33_NO_~~4~~_ | 399.53 | 400.2460 | -6.74 | 382.2382, 372.2541, 340.1917, 332.1856, 314.1768, 290.1384, 288.1589, 274.1441, 271.2061, 264.1235, 243.2107, 215.1797, 189.1642, 175.1486, 173.1332, 156.0295, 147.1171, 142.0501, 138.0186, 137.1325, 130.0502, 121.1013, 109.1012, 107.0856, 95.0855, 81.0699, 69.0700, 67.0544, 60.0445, 55.0545 | - | + |
| 23* | (E)-3-(hydroxy(1,3,6-trimethyl-2-((E)-prop-1-en-1-yl)-1,2,4a,5,6,7,8,8a-octahydronaphthalen-1-yl)methylene)-5-(hydroxymethyl)pyrrolidine-2,4-dione | 20.03 | 374.2331 | [M+H]^+^ | C_22_H_31_NO_4_ | 373.49 | 374.2327 | -1.06 | 356.2219, 346.2376, 328.2273, 304.1536, 245.1098, 217.1953, 191.1796, 170.0813, 156.0656, 95.0854 | + | + |
| 24* | (E)-3-((1,6-dimethyl-2-((E)-prop-1-en-1-yl)-1,2,4a,5,6,7,8,8a-octahydronaphthalen-1-yl)(hydroxy)methylene)-5-(hydroxymethyl)-1-methylpyrrolidine-2,4-dione/ equisetin | 19.68 | 374.2331 | [M+H]^+^ | C_22_H_31_NO_4_ | 373.49 | 374.2324 | -1.87 | 356.2224, 346.2379,  288.1584, 252.1229, 170.0814, 144.0657, 126.0554, 95.0856 | + | + |
| 25* | (Z)-3-(hydroxy(1,3,6-trimethyl-2-((E)-prop-1-en-1-yl)-1,4,4a,5,6,7,8,8a-octahydronaphthalen-1-yl)methylene)-1-methyl-1,3-dihydro-2H-4l5-pyrrole-2,4-dione | 21.27 | 358.2382 | [M+H]^+^ | C_22_H_31_NO_3_ | 357.49 | 358.2376 | -1.67 | 340.2277, 330.2433, 217.1955, 191.1798, 140.0708, 95.0855 | - | + |
| 26* | (E)-3-(hydroxy(1,3,6-trimethyl-2-((E)-prop-1-en-1-yl)-1,2,4a,5,6,7,8,8a-octahydronaphthalen-1-yl)methylene)-1-methyl-5-methylenepyrrolidine-2,4-dione | 20.55 | 370.2387 | [M+H]^+^ | C_23_H_31_NO_3_ | 369.51 | 370.2381 | -1.62 | 342.2430, 340.2266, 288.1600, 227.1798, 191.1797, 152.0708, 95.0855 | - | + |

#### Table S6. MIC values of compound 1 against ESKAPE strains.

| **Bacterial strain** | **MIC (µg/mL)** |
| --- | --- |
| **Gram negative** |  |
| E. cloacae ATCC 13047 | > 64 |
| K. pneumoniae ATCC 138833 | > 64 |
| A. baumannii ATCC 17978 | > 64 |
| P. aeruginosa ATCC 27853 | > 64 |
| E. coli ATCC 25922 | > 64 |
| **Gram positive** |  |
| E. faecalis ATCC 29212 | 4 |
| S. aureus ATCC 29213 | 1 |
| S. aureus NCTC 8325 | 1 |
| S. aureus Newman | 1 |
| S. aureus HG001 | 2 |

### Supplementary Figures

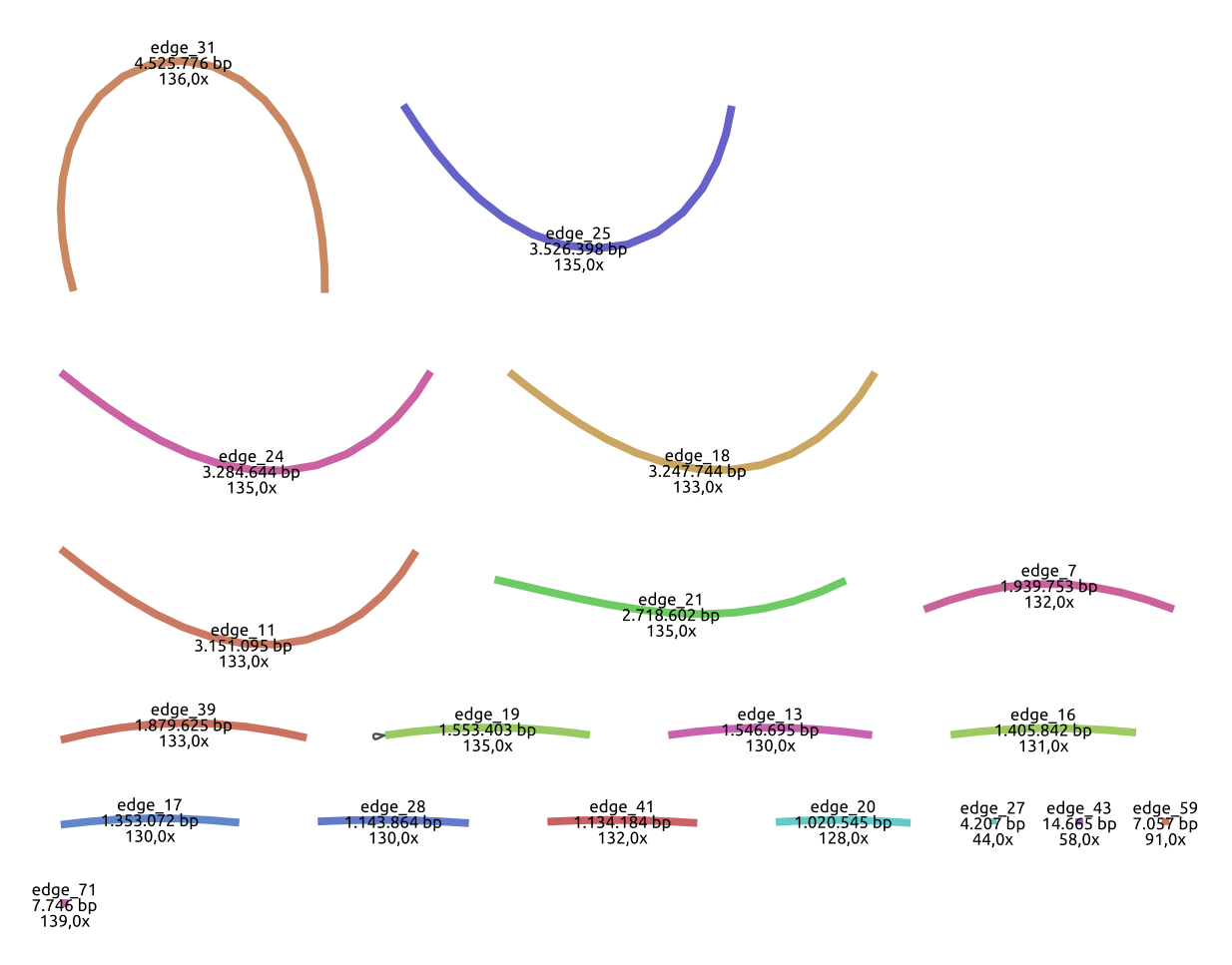

#### Figure S1. Contigs of *Neocucurbitaria* sp. VM-36 visualized by Bandage.

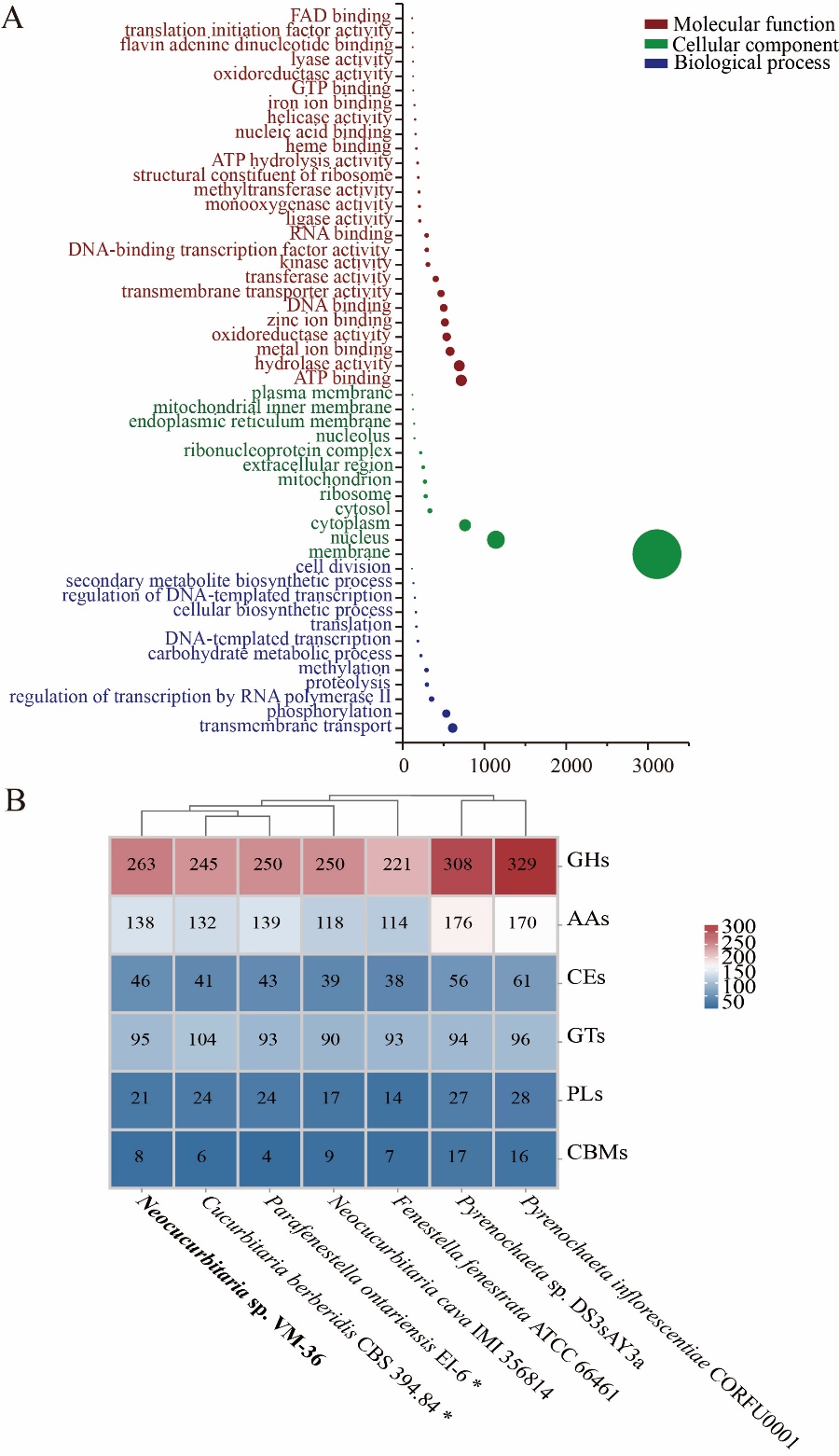

Figure S2. Functional genome annotation of *Neocucurbitaria* sp. VM-36. (A) Gene Ontology (GO) annotation (Top 50 GO terms). The size of each dot corresponds to the number of genes enriched in a given GO term compared to the background gene set. (B) CAZyme analysis and comparison to other *Cucurbitariaceae* species; GHs, glycoside hydrolases; AAs, auxiliary activities; CEs, carbohydrate esterase; GTs, glycosyl transferases; PLs, polysaccharide lyases; CBMs, carbohydrate binding modules. Strains marked with an asterisk (*) represent Type strains.

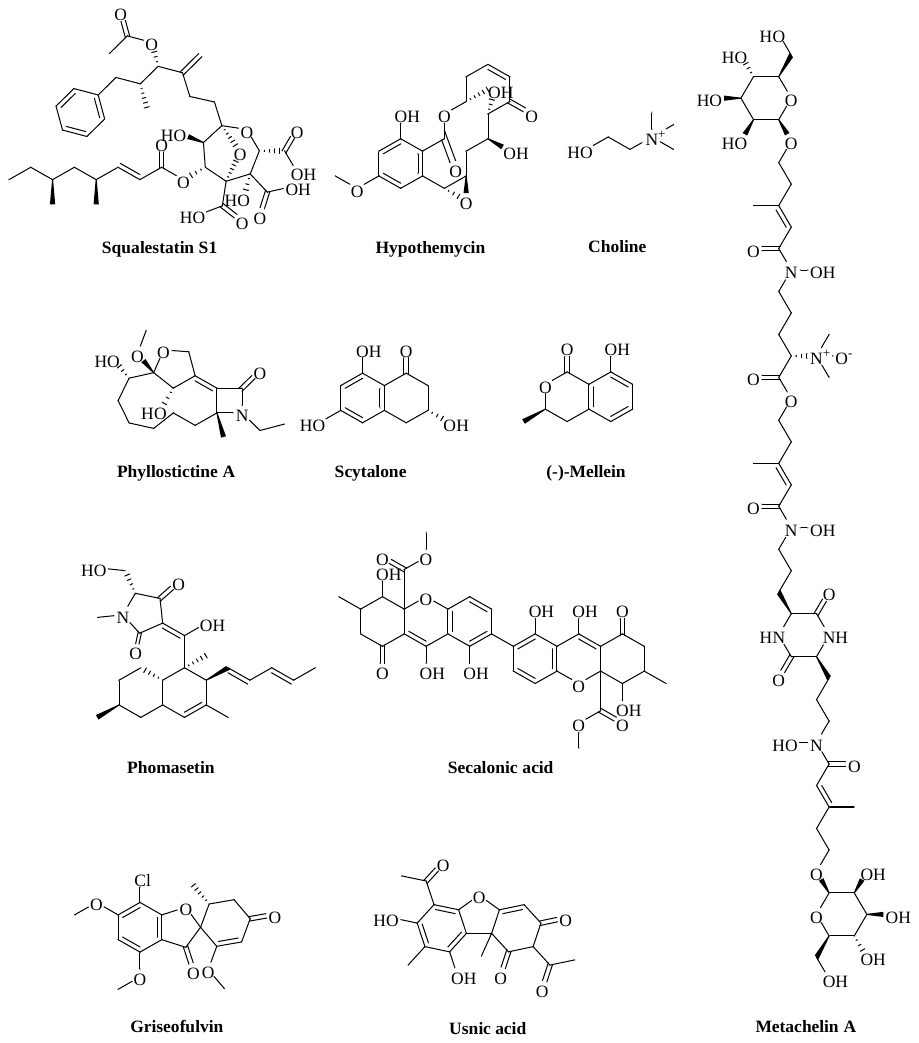

#### Figure S3. Structures of compounds predicted to be produced by *Neocucurbitaria* sp. VM-36 based on AntiSMASH analysis.

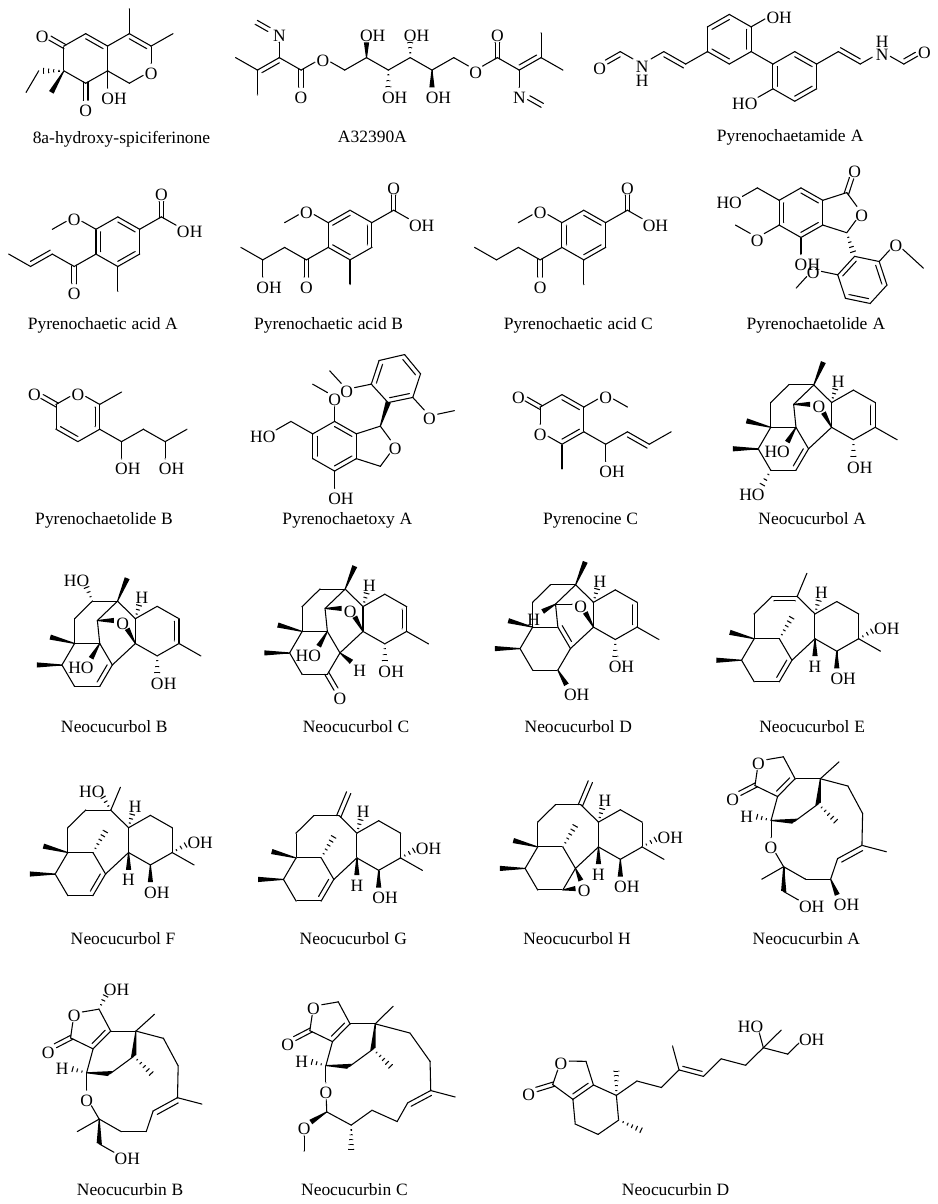

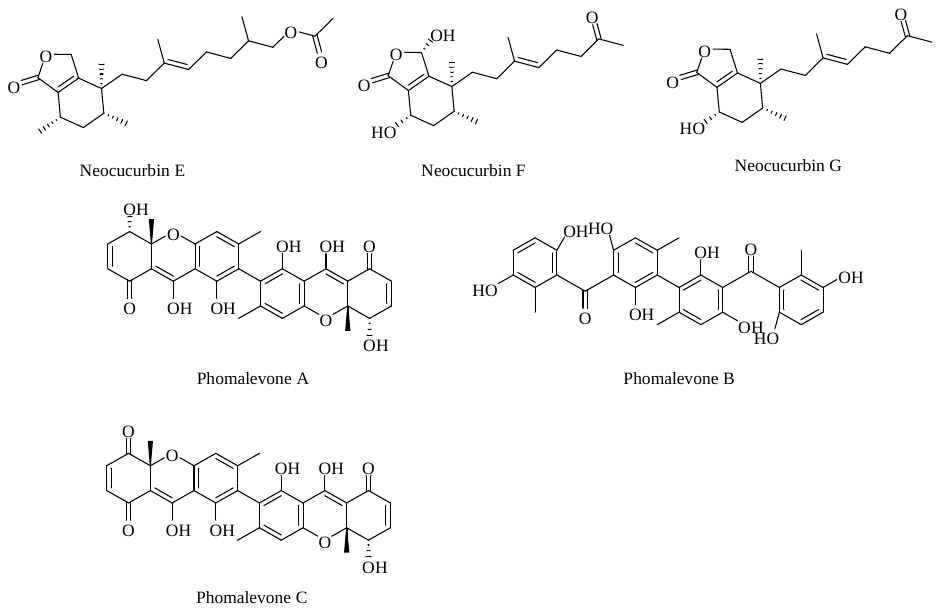

#### Figure S4. Structures of compounds reported to be produced by fungi from the *Cucurbitariaceae* family. See Table S4 for literature related to these compounds.

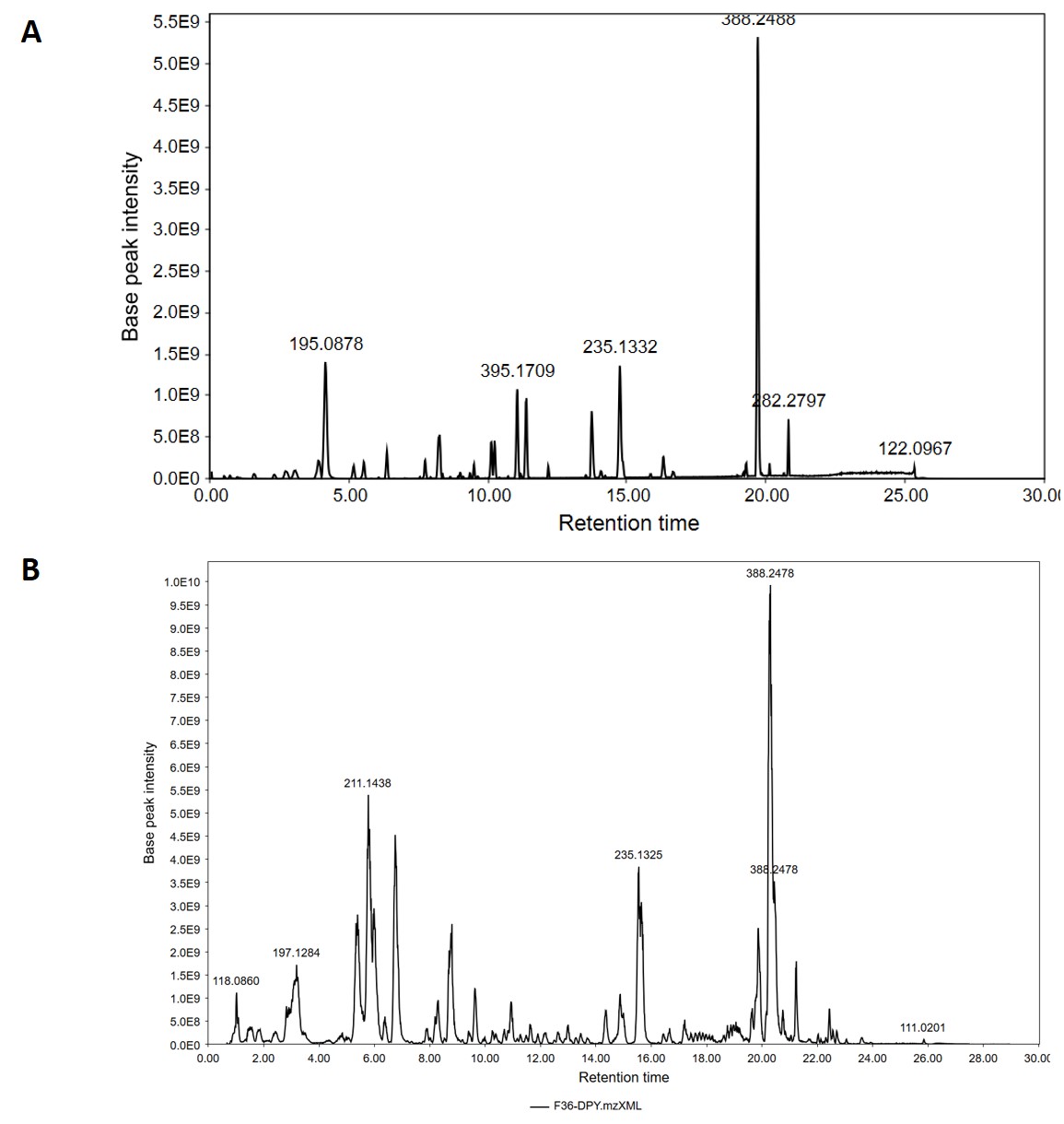

#### Figure S5. Total ion chromatograms recorded in positive ion mode of EtOAc extracts of *Neocucurbitaria* sp. VM-36 grown on PDA solid medium and DPY liquid medium. (A) *Neocucurbitaria* sp. VM-36 grown on the PDA solid medium at 25 ℃ for 28 days; (B) *Neocucurbitaria* sp. VM-36 grown in the DPY liquid medium at 25 ℃ for 30 days, with shaking at 130 rpm.

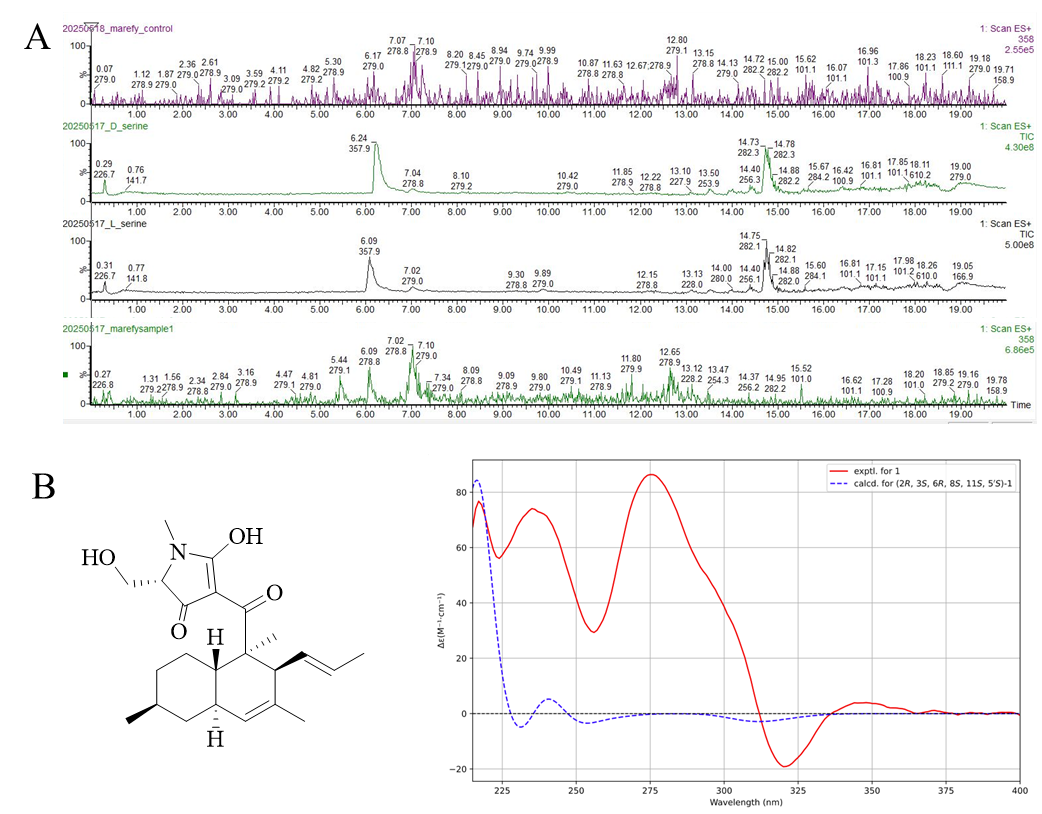

#### Figure S6. Structural characterization of compound 1 and its isomer.

Marfey’s result of compound **1** compared with standards L-serine and D-serine (A). Structure and calculated ECD spectra of the compound **1** isomer with C-1 being a carbonyl group and C-2’ being a hydroxyl group (B).

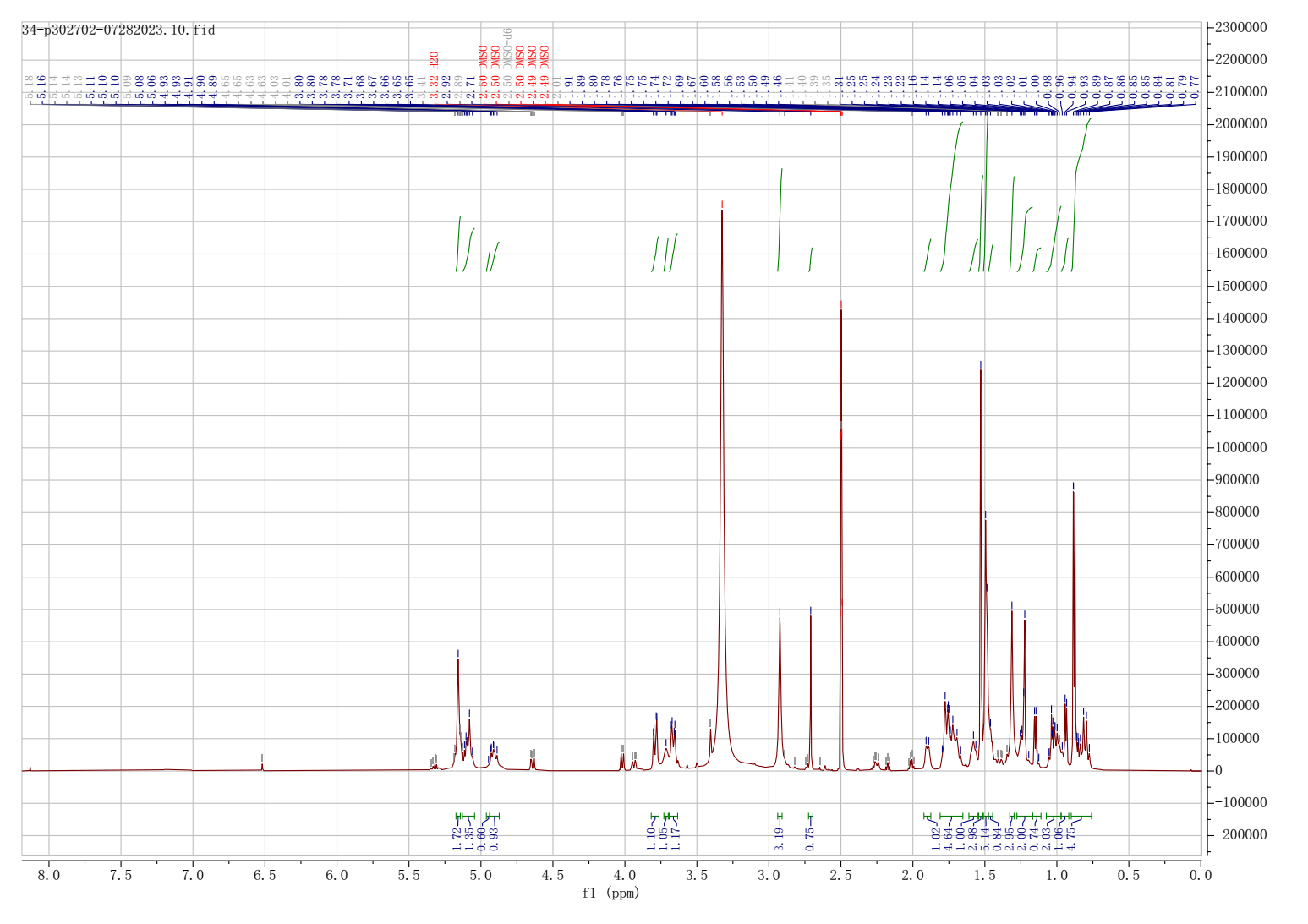

#### Figure S7. ^1^H NMR spectrum of compound 1 in DMSO-d6 (600 MHz).

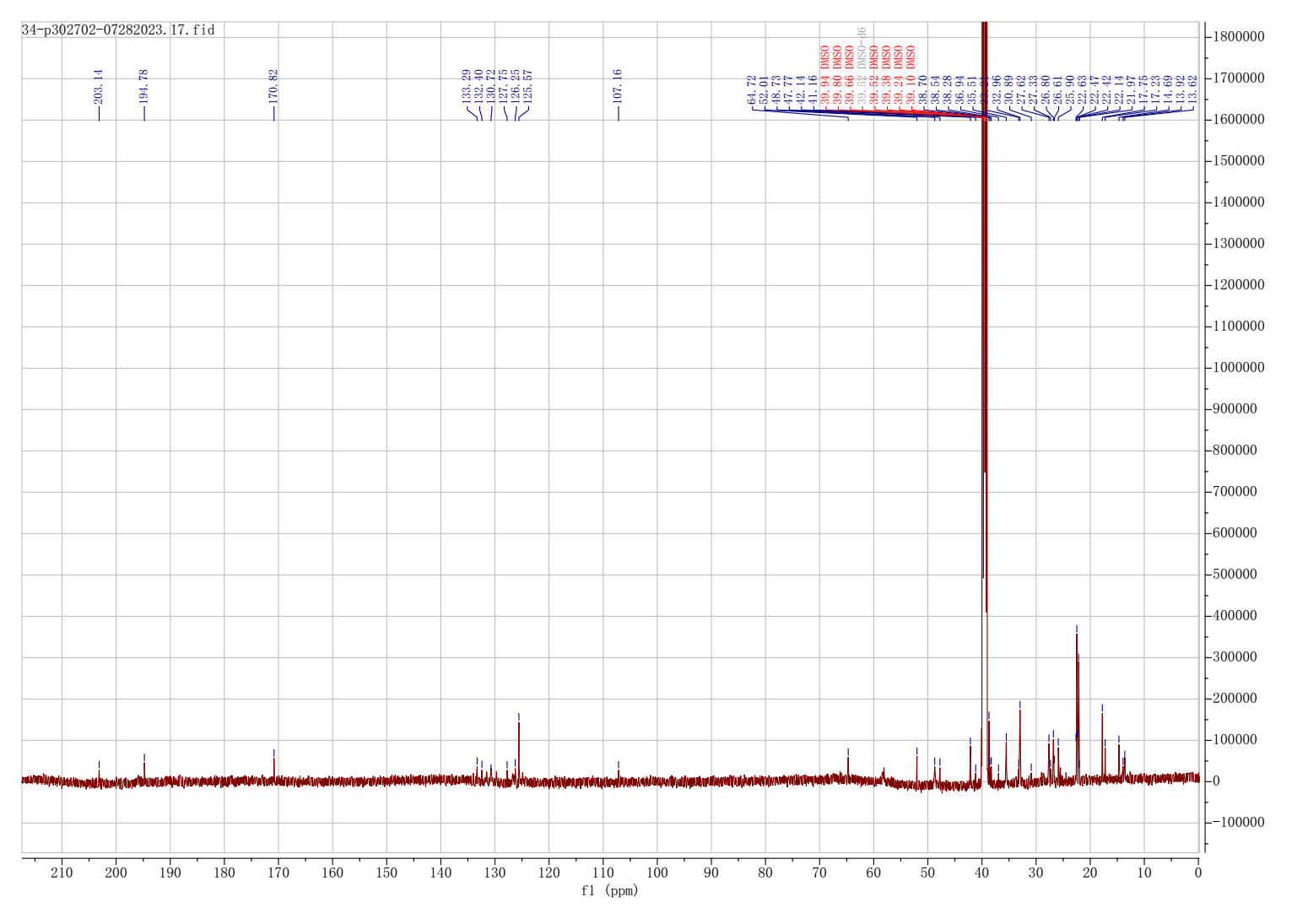

#### Figure S8. ^13^C NMR spectrum of compound 1 in DMSO-d6 (150 MHz).

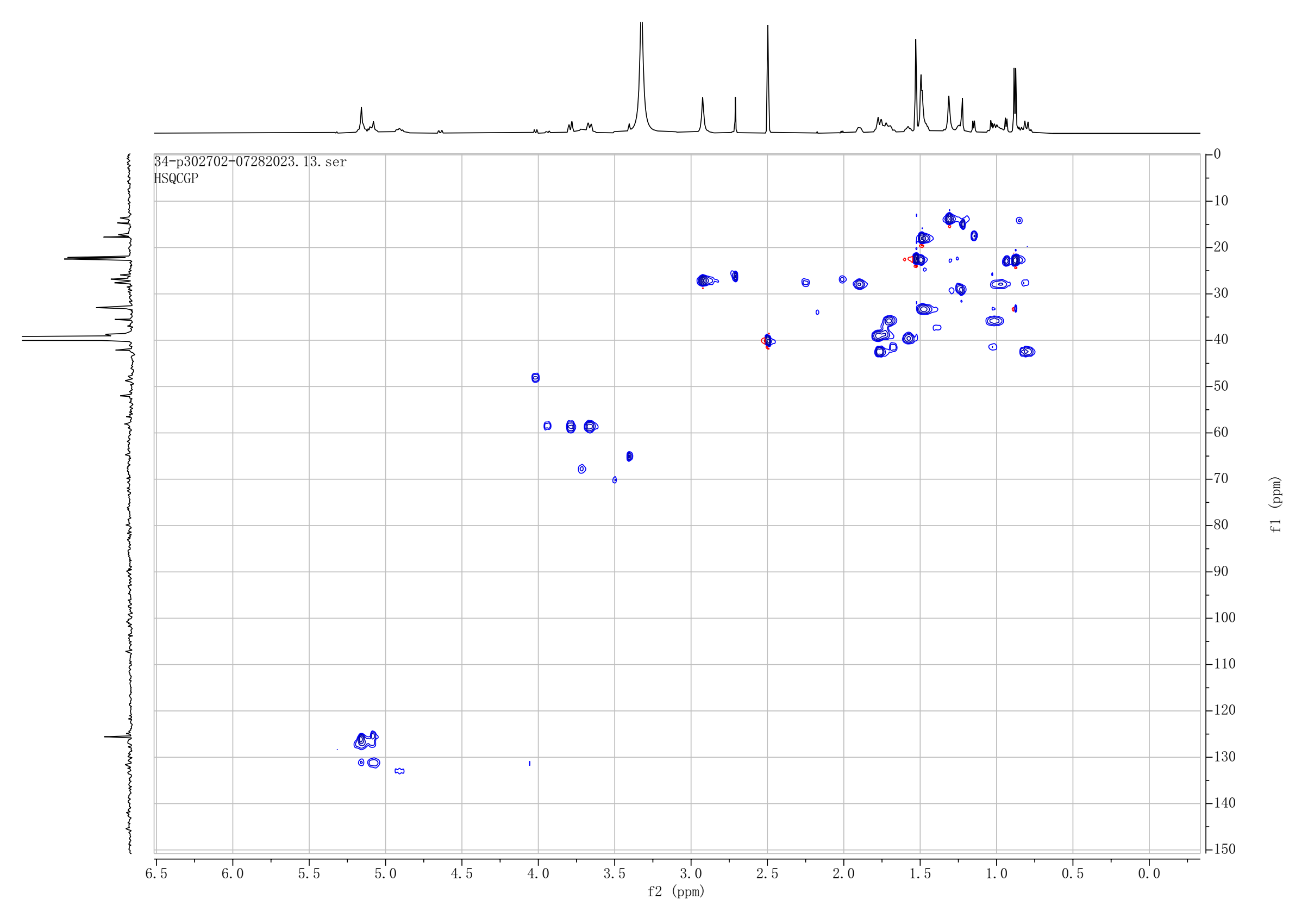

#### Figure S9. HSQC spectrum of compound 1 in DMSO-d6.

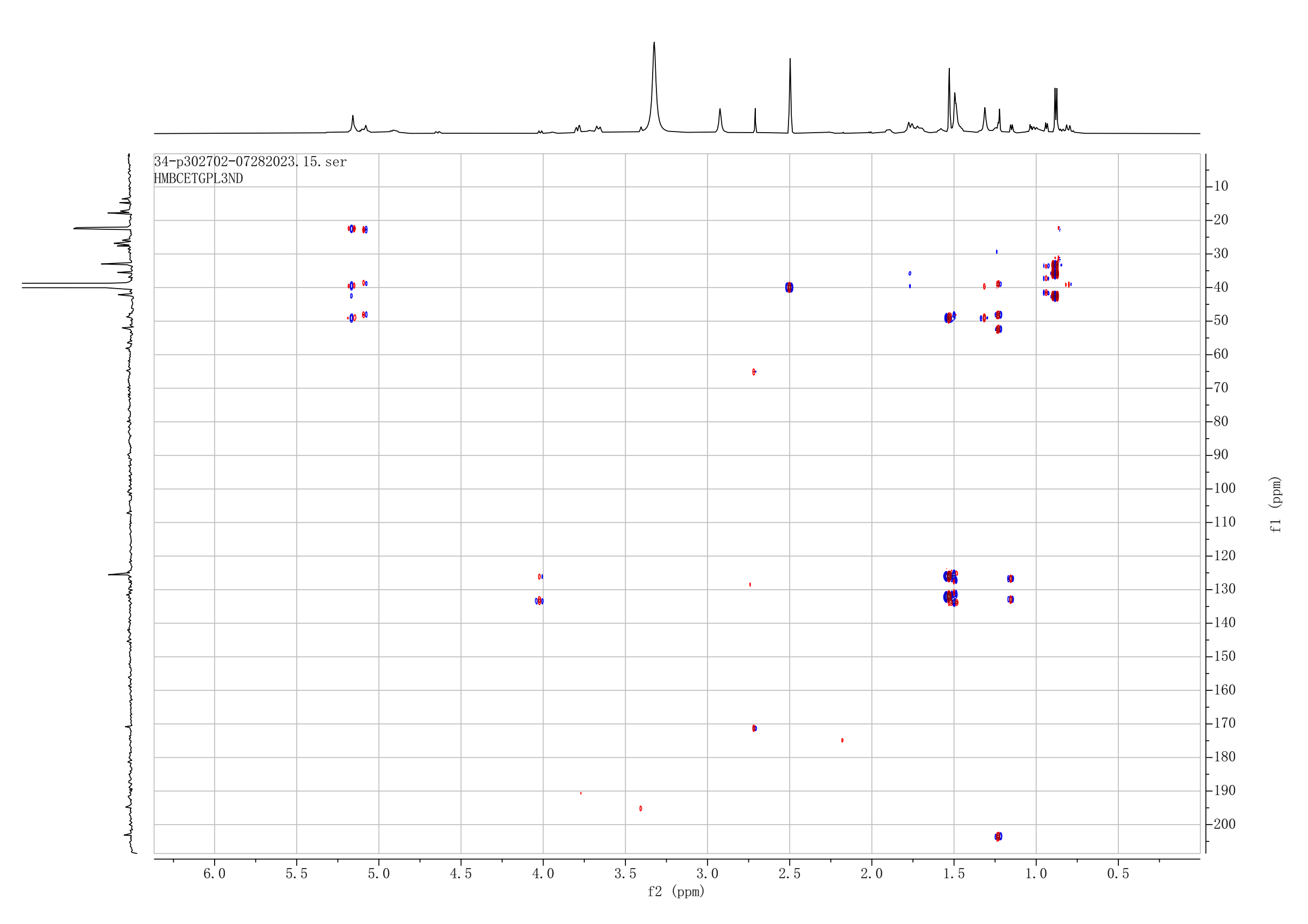

#### Figure S10. HMBC spectrum of compound 1 in DMSO-d6.

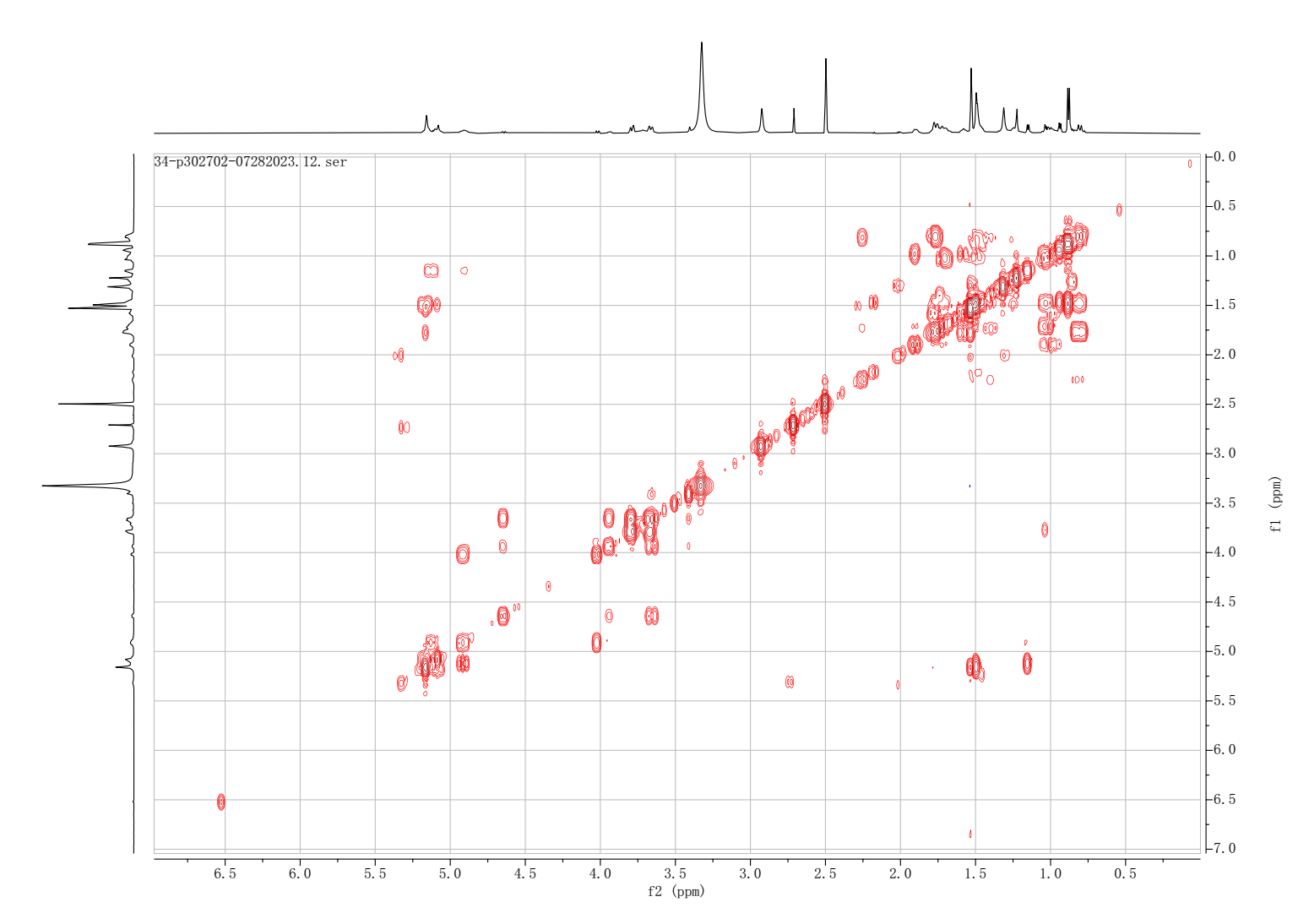

#### Figure S11. ^1^H-^1^H COSY spectrum of compound 1 in DMSO-d6.

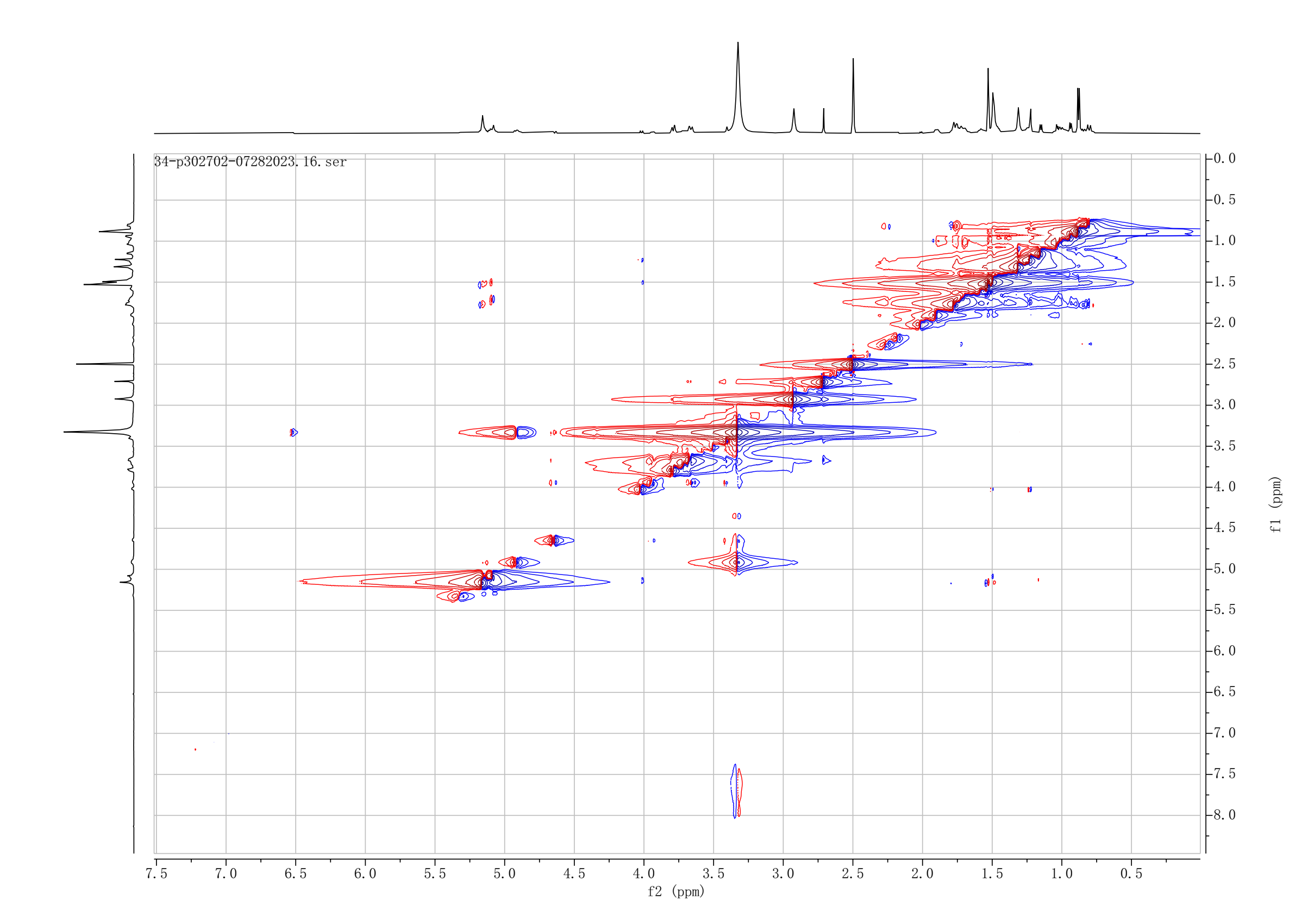

Figure S12. NOESY spectrum of compound 1 in DMSO-d6.
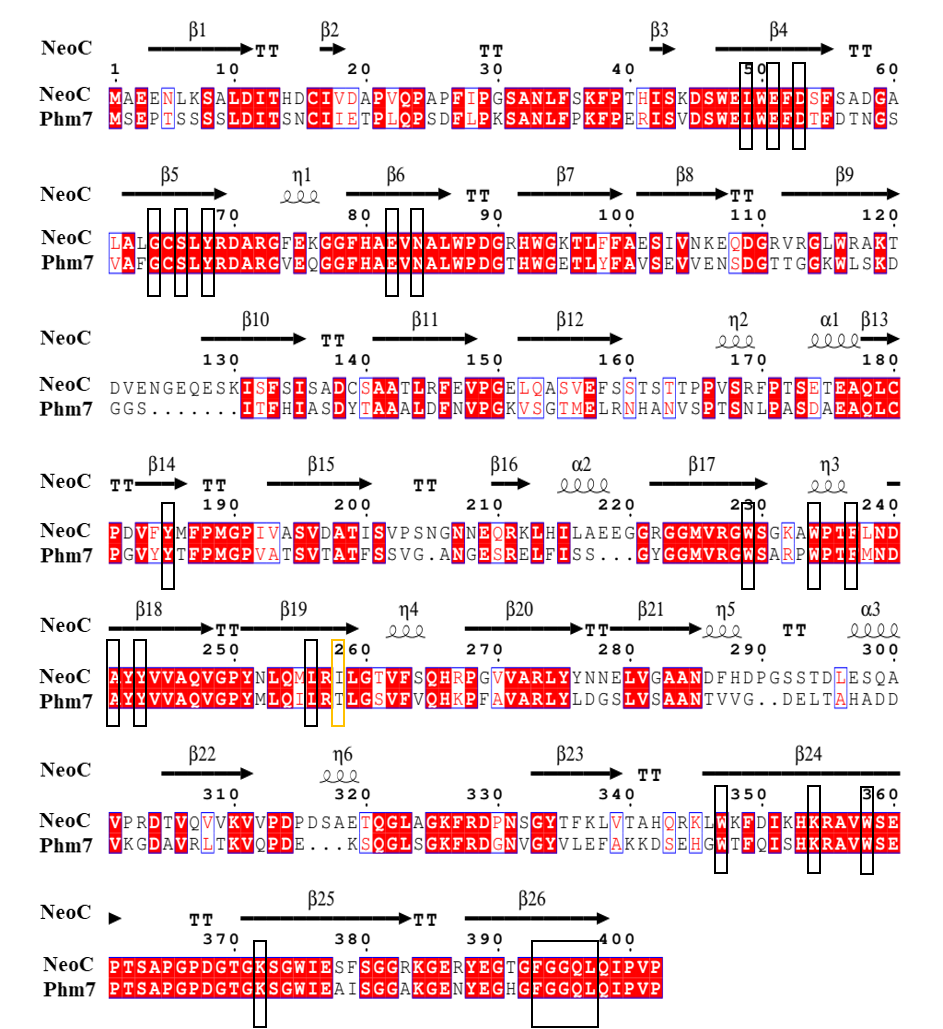

#### Figure S13. Amino acid sequence alignment between NeoC of *Neocucurbitaria* sp. VM-36 and Phm7 of *Pyrenochaetopsis* sp. RK10-F058.

The alignment was performed with ClustalW ^48^ and displayed in ESPript 3.0 ^49^. The Phm7 sequence was obtained from the MiBIG database (protein BBC43190.1). In the ESPript 3.0 output, fully conserved residues are shown as white letters on a red background. Red letters with small blue boxes indicate highly similar residues with conserved physicochemical properties. Residues without color or framing are variable and less conserved. Amino acids inside the large black boxes were previously implicated in catalytic activity ^2,35,50^. The yellow box highlights amino acid 258 where isoleucine in NeoC replaces threonine in Phm7.

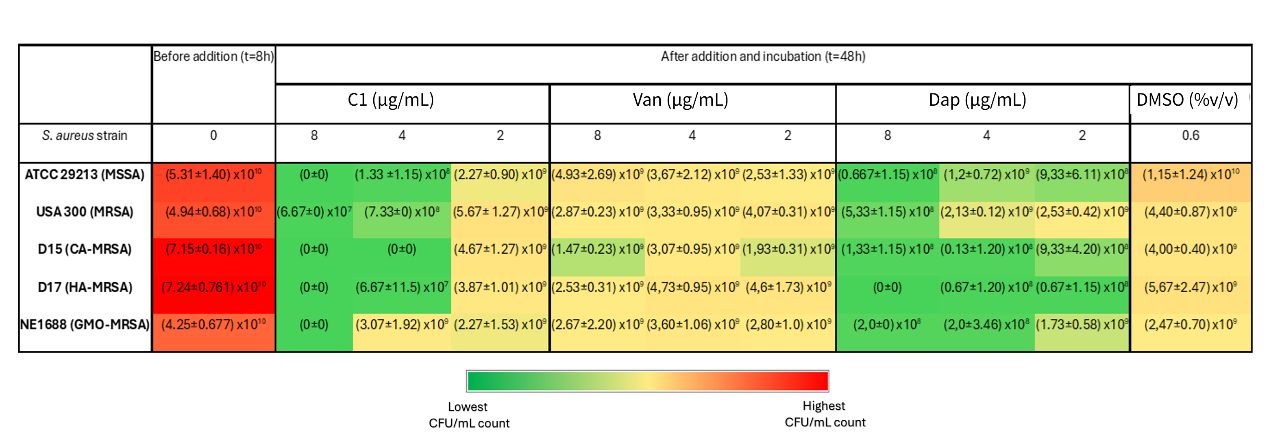

#### Figure S14. Bacterial viability analysis before (t = 8 h) and after the addition of compound 1, vancomycin or daptomycin and incubation (t = 48 h). The heatmap represents mean values of CFU/mL ± standard deviation (n = 3) with the colour code being based on the mean values.

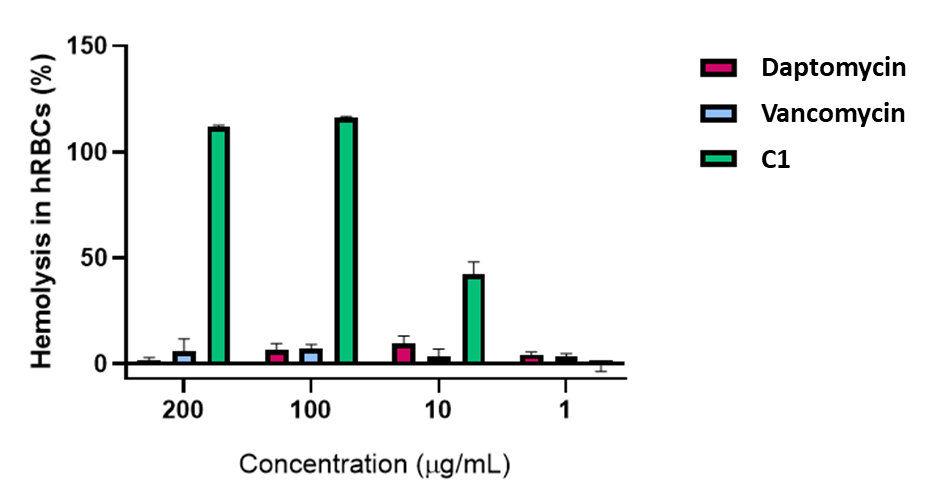

#### Figure S15. In vitro hemolytic activity of compound 1 towards human red blood cells as compared to vancomycin and daptomycin.
